# Genome-scale label-free imaging reveals cellular physiology encoded in bacterial collective architecture

**DOI:** 10.64898/2026.08.30.748126

**Authors:** Seh Na S. Mellick, Jesse J. Derringer, Darin Boyes, Gillian Croteau, Madeline Burke, Steven Gifford, Drew J. Stark, Laura A. Mike, Sophia Turecki, Oana Carja, Irina V. Mikheyeva, Andrew A. Bridges

## Abstract

DNA sequencing unified microbial genotyping into a single, comprehensive readout, yet phenotyping remains a slow and fragmented endeavor. Here, we introduce **Micro**bial **P**henotyping **U**sing **L**ow-magnification **L**abel-free **I**maging (µPULLI), a computer vision platform that extracts microcolony and population-level phenotypes from brightfield timelapses of liquid culture growth. Using µPULLI, we screened a genome-scale *Vibrio cholerae* mutant library, recording more than 200,000 images, which revealed that core bacterial pathways shape community architecture. Functionally related mutants converge in appearance, allowing us to resolve processes as distinct as biofilm formation, motility, central metabolism, cofactor biosynthesis, and envelope composition using a single approach. We further show µPULLI can be used to determine a drug target, characterize other pathogens, and classify bacterial species. Our results show that bacterial multicellular development is an interpretable signature of genotype-phenotype relationships, which can be captured from simple brightfield timelapses. We release the µPULLI pipeline and an interactive atlas of community forms.

## INTRODUCTION

A fundamental question in biology is how the physiological state of single cells shapes the architecture of the communities they build. In multicellular organisms the link is well established: tissue architecture reflects not only the activity of dedicated patterning genes but the integrated state of core cellular processes. Disrupting mitochondrial function can reshape tissue form as profoundly as mutating a canonical developmental regulator.^1,2^ Bacteria, though unicellular, likewise organize into emergent multicellular collectives (e.g. biofilms, aggregates, and swarms) whose architecture emerges from the coordinated behavior of individual cells.^3^ Thus, in microbes the same logic should hold, such that disrupting core metabolic and physiological processes at the cellular level should manifest in reproducibly altered multicellular, community-level phenotypes. Individual links between physiological pathways and bacterial collective behavior have been extensively reported,^4-6^ but the relationship between cell physiology and community morphology has never been characterized systematically. Assessing the extent to which cellular physiology influences collective form therefore requires genome-scale phenotyping methods that capture a comprehensive view of community development and do not presuppose which pathways are of importance.

Over the past decade, DNA sequencing technologies have revolutionized researchers’ abilities to genotype the microbial world due to declining costs and widespread availability.^7^ In contrast, behavioral phenotyping has not kept pace. Microbial collective behaviors such as surface attachment, motility, and biofilm formation, which govern how bacteria colonize surfaces, resist clearance, and persist during infection, are typically evaluated using separate assays,^8,9^ each optimized for measuring a single trait at a single timepoint. Moreover, each assay can only report the phenotypic trait it was designed to measure, so what can be discovered is fixed in advance to phenotypes for which an assay and vocabulary already exist. As such, informative phenotypes that fall outside of expectations often go unclassified. This fragmentation limits the ability of researchers to capture a comprehensive view of bacterial behavior and hinders the systematic discovery of shared or multifunctional regulatory pathways.

Recent advances in microscopy, computer vision, and machine learning have begun to expand the scope of microbial phenotyping, enabling quantitative interrogation of colony morphology and cell shapes.^10,11^ However, these approaches generally remain narrowly tailored to specific phenotypes, prioritize predictive performance over mechanistic interpretability, or depend on fluorescence labeling and high-resolution imaging to resolve single-cell behaviors.^12-14^ Thus, an approach is needed to characterize microbial behavior broadly and interpretably without presupposing which phenotypes matter, and that is readily scalable to enable phenotype characterization across large microbial libraries.

Here we introduce µPULLI (<u>Micro</u>bial <u>P</u>henotyping <u>U</u>sing <u>L</u>ow-magnification <u>L</u>abel-free Imaging), a unified platform that characterizes microcolony and population-level bacterial behaviors from brightfield timelapses of cultures growing in 96-well plates. Image acquisition requires a standard inverted microscope and a 10× air objective, and does not require stains, reporters, or specialized growth conditions. We provide two analysis routes: µPULLI-DL, which passes images through a deep-learning (DL) vision transformer-based analysis pipeline to extract image embeddings without fine-tuning, and µPULLI-I, which extracts interpretable (I) biofilm biomass, microcolony, and texture features that can be traced back to visible structures within the images. Applying µPULLI to an ordered *Vibrio cholerae* transposon library,^15-17^ we recorded more than 200,000 images and assembled an atlas describing variation in community development and morphology. µPULLI revealed that perturbations to central metabolism, cofactor biosynthesis, nutrient acquisition, and cell-envelope composition each left reproducible, pathway-specific marks on multicellular developmental dynamics, and functionally related mutants converged on similar community-level phenotypes. A single µPULLI experiment therefore resolves diverse pathways and phenotypes, each of which would conventionally require its own assay. We also showed that pharmacological inhibition of a core biotin biosynthesis enzyme phenocopied deletion of genes involved in biotin biosynthesis, demonstrating that drug-candidates can be assigned agnostically to their target pathway from images alone. The approach extended without modification to classifying *Klebsiella pneumoniae* mutants based on multicellular phenotype, and to species classification across eight taxonomically diverse organisms, demonstrating the broad utility of µPULLI. We release the atlas, interactive viewers of the phenotype landscape, the analysis code, and the robotic liquid-handling protocols, so that any laboratory with an inverted microscope can generate comparable data and project new strains, species, or compounds onto the same coordinate system.

## RESULTS

### Low magnification timelapses of liquid cultures carry genotype-specific information

For decades, optical density (OD) measurements have served as the standard for monitoring microbial growth. However, this bulk approach reduces the spatial and temporal heterogeneity of a growing culture to a single scalar readout. We hypothesized that low-magnification brightfield timelapse imaging of microbial growth in liquid culture would capture spatially resolved community phenotypes that OD cannot (Fig. 1A). Importantly, the approach requires no specialized growth or imaging; timelapse brightfield images of static bacterial cultures growing in 96-well plates were captured using a 10× air objective. To assess whether such images carry genotype-specific information, we assembled a panel of eight previously characterized *V. cholerae* strains: the WT parent; three mutants with altered biofilm matrix machinery, Δ*vpsL* (matrix deficient),^18^ *vpvC^W240R^* (hyper-matrix producer),^19^ and Δ*rbmB* (matrix degradation deficient);^20,21^ and four mutants in pathways adjacent to the biofilm lifecycle, covering motility (Δ*flaA*),^22^ quorum sensing (phenotypically distinct mutants Δ*hapR* and *luxO^D61E^*),^23^ and polyamine import (Δ*potD1*) (Fig. 1B).^24^ Consistent with previous results, WT *V. cholerae* transiently assembled into microcolony biofilms that dispersed as the culture matured (Fig. 1B, Fig. S1, Movie S1).^25,26^ Against that baseline, every mutant we examined displayed altered spatiotemporal dynamics with distinctive visual characteristics (Fig. 1B, Fig. S1A, Movie S1).

**Figure 1.**
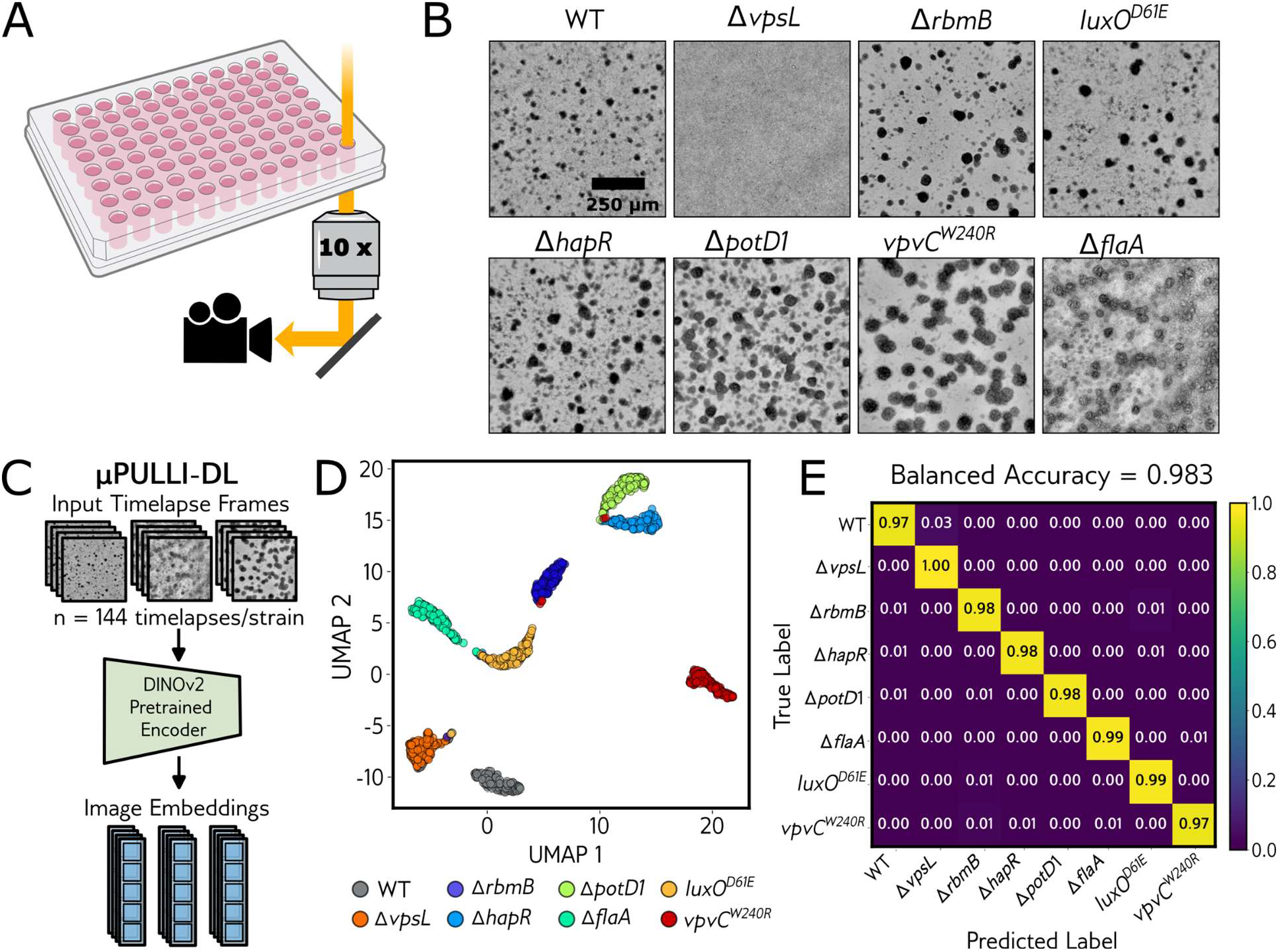
µPULLI-DL differentiates genotypes from low-magnification brightfield timelapses. **(A)** Schematic of data acquisition. Static bacterial growth in 96-well plates was captured at 1-hour intervals for 30 hours via brightfield microscopy using a 10× air objective lens. **(B)** Representative low-magnification images of the indicated *V. cholerae* strains, each shown at their peak biofilm biomass timepoint. Temporal dynamics can be viewed in Fig. S1, Movie S1. **(C)** Schematic of the computer vision pipeline; 144 videos per strain were preprocessed and then passed through a DINOv2 vision transformer (ViT-B/14) without fine-tuning to extract per-frame classification (CLS) token image embeddings. **(D)** Dimensionality reduction of the 768-dimensional DINOv2 CLS embeddings stacked across 31 timepoints (0-30 h) using UMAP (n_neighbors = 10, min_dist = 0.1), where point colors denote distinct *V. cholerae* strains. Each point represents a single replicate timelapse video, n ≈ 144 replicate videos per strain. **(E)** Confusion matrix for a random-forest classifier predicting the eight *V. cholerae* genotypes from stacked DINOv2 CLS embeddings (using timepoints spanning the growth window, 9–30 hours). Evaluation used plate-held-out cross-validation, in which replicates from the same plate were never split across training and test sets (see Methods). Rows are the true genotype, columns are the predicted genotype; color is the row-normalized fraction of held-out wells (0–1; diagonal = per-genotype recall), averaged over folds. Mean balanced accuracy = 98.3%. *N* = 1,149 wells (approximately 144 per genotype) imaged across 18 plates. a.u., arbitrary units.

To test whether such qualitative distinctions between strains are reproducible and quantifiable, we used robotic liquid handling to generate replicate timelapses of each strain for deep learning feature extraction (144 replicates per strain). Each timelapse was passed through the pre-trained encoder layer of DINOv2,^27^ a vision transformer model pretrained on natural images from the web and applied here without fine-tuning (Fig. 1C), allowing us to test whether genotype-specific signal is recoverable from low-magnification brightfield imaging without any prior assumptions about which visual characteristics are of importance. Consistent with our qualitative observations, DINOv2 embeddings for strain replicates grouped by genotype in UMAP space^28^ (Fig. 1D, Fig. S1B), and, remarkably, a random-forest classifier trained on embedding trajectories distinguished all eight strains at 98.3% balanced classification accuracy (Fig. 1E, Fig. S1C). Thus, genotypes are reproducibly encoded within the visible spatiotemporal dynamics of culture growth, and recoverable by a model that was never trained on biological or medical images. Whereas resolving matrix production, motility, quorum sensing, and polyamine transport would conventionally require a separate assay for measuring each characteristic, our simple approach distinguishes perturbations to these pathways using a single label-free imaging assay. We refer to this approach as <u>M</u>icrobial <u>P</u>henotyping <u>U</u>sing <u>L</u>ow-magnification <u>L</u>abel-free Imaging (µPULLI), and we designate the deep-learning vision transformer-based analysis used here as µPULLI-DL.

### Quantitative feature extraction for interpretable genotype classification

Although the DINOv2 embeddings achieved highly accurate genotype classification, they provide no insight into which visual properties distinguish genotypes. We therefore turned to classical image analysis, extracting explicitly defined, human-interpretable measurements from the same images (Fig. 2A). Previously, we developed a framework to quantify overall microcolony biofilm biomass within the field of view from low-magnification brightfield timelapse videos.^29^ Applying this approach to the eight previously characterized *V. cholerae* strains revealed highly reproducible, strain-specific biofilm trajectories. Notably, the strains differed in both the magnitude of microcolony formation and the rate of dispersal (Fig. 2B). As such, we asked whether biofilm biomass trajectories alone could distinguish strains. Indeed, a random-forest classifier trained on biofilm biomass trajectories achieved 93.8% balanced accuracy across the eight strains, approaching the classification accuracy obtained using DINOv2 embeddings (Fig. S2A). However, biofilm biomass collapses each frame to a single scalar value, discarding how colonies are shaped, arranged, and textured. We therefore developed a microcolony-tracking method to track and segment *V. cholerae* microcolony biofilms as they form and disperse, allowing us to expand our feature set beyond the single biofilm biomass metric to include 12 colony segmentation-derived features describing the shape, intensity, and spatial arrangement of individual segmented microcolony biofilms. In addition, we measured 14 whole-image texture features, 13 of which are Haralick textural statistics^30^ quantifying spatial relationships between pixel intensities (Fig. 2C, see Table S1 for a description of each feature class). Supplementing the biofilm biomass metric with the 26 colony-level and whole-image features raised balanced classification accuracy from 93.8% to 99.0%, comparable to the classification performance of µPULLI-DL (98.3%), but now built entirely from interpretable, physically defined measurements, with replicates of different genotypes separating almost completely in UMAP (Fig. 2D, E). We refer to this interpretable implementation as µPULLI-Interpretable (µPULLI-I).

**Figure 2.**
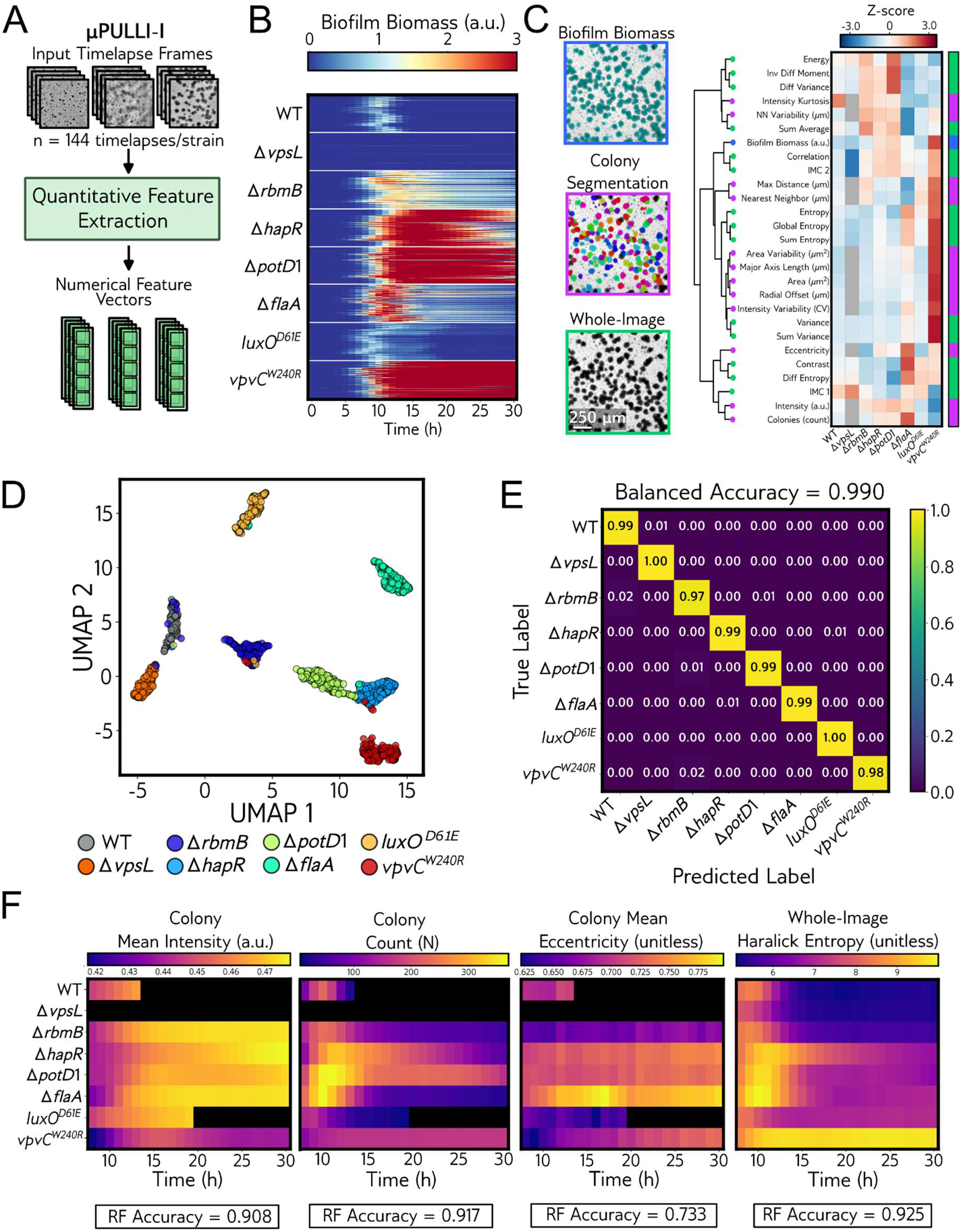
µPULLI-I classifies genotypes from interpretable quantitative image features. **(A)** Schematic of data acquisition and feature extraction. Image data are the same as in Fig. 1. Here, we used quantitative feature extraction rather than deep-learning embeddings, yielding metrics for bulk biofilm biomass, colony-level features, and whole-image statistics per frame. **(B)** Heatmap illustrating whole field of view biofilm biomass trajectories for each *V. cholerae* strain. All values are normalized to the mean peak biofilm biomass of the WT strain. Each horizontal trace represents an individual timelapse replicate. **(C)** Left: Representative images showcasing the three categories of quantitative features extracted in this study. Biofilm biomass (blue, top panel), computed through a simple threshold on the intensity values of the processed images, encompasses the light scattered by biofilm regions in the field of view; colony segmentation-derived features (magenta, middle panel) capture the spatial and optical properties of individual segmented colonies, averaged over all colonies in the well; and whole-image features (green, bottom panel) capture image texture statistics extracted from unlabeled processed images. Right: Heatmap of Z-scores for the indicated features at each strain’s peak biofilm biomass timepoint, calculated across all strains from the median of 144 replicates per strain. Rows (features) are ordered by hierarchical clustering (Ward linkage, Euclidean distance); strain order is fixed. Gray denotes absence of colony-level features. Colorbar denotes the class of features being measured. **(D)** Dimensionality reduction using UMAP visualizes all measured features for each timelapse replicate, with point colors indicating the corresponding *V. cholerae* strains. **(E)** Confusion matrix for a random-forest classifier predicting the eight *V. cholerae* genotypes from quantitative texture and morphological features, evaluated by plate-held-out cross-validation (see Methods). Rows are the true genotype, columns are the predicted genotype; color is the row-normalized fraction of held-out wells (0–1; diagonal = per-genotype recall), averaged over folds. Mean balanced accuracy = 99.0%. *N* = 1,149 wells, approximately 144 wells per genotype, imaged across 18 plates. **(F)** Heatmaps of average trajectories for the indicated feature classes for each *V. cholerae* strain (*N* = 144). Balanced random forest (RF) classification accuracy for each isolated feature class is shown below the plot. Temporal heatmap plots for all feature classes are provided in Fig. S3. Black denotes absence of colony-level features. a.u., arbitrary units.

Unlike the DINOv2 embeddings, µPULLI-I let us ask which visual properties distinguish genotypes. Classifying the eight strains on each feature class independently, we found that colony segmentation-derived features and whole-image texture features yielded balanced classification accuracies of 96.0% and 98.5%, respectively, approaching the accuracy achieved on the full feature set (Fig. S2B, S2C). Either feature class alone was sufficient for discriminating strain identity, indicating that genotype is redundantly encoded in spatial and geometric properties of single colonies, as well as whole-image texture. We retained the colony-level features despite relatively lower classification performance (<3% difference in balanced classification accuracy) because, unlike whole-image texture statistics which do not require accurate segmentation, they correspond to interpretable, physical properties of microcolony biofilms. Among colony-level feature trajectories, mean colony intensity (90.8% classification accuracy) and colony count (91.7%) were the strongest predictors of genotype when used independently for classification, while colony radial offset (88.1%), colony area (82.6%), and eccentricity (73.3%) were weaker but still informative (Fig. 2F, Fig. S2D, and Fig. S3). Among whole-image feature trajectories, a single texture statistic, Haralick sum entropy, reached 97.3% balanced classification accuracy when used independently (Fig. S2D, Fig. S3), nearly matching the accuracy obtained on the full feature set and underscoring the power of segmentation-free texture statistics for quantifying the emergent spatial and textural patterns that distinguish these genotypes (for more interpretation of Haralick textural statistics, see Supplementary Text). Combining the biofilm biomass, colony-level, and whole-image textural features therefore yields a classifier that is both interpretable and broadly applicable, capturing genotype-specific signals regardless of whether individual microcolonies can be resolved.

Because microcolonies form, grow, and disperse over the course of the experiment, we next asked when genotypes become distinguishable during this progression. Training a separate classifier on the full feature set at each timepoint, we found accuracy near chance for the first five hours post-inoculation, before microcolonies became visible, then rising steadily to ∼96% balanced classification accuracy by ∼17 hours and plateauing thereafter (Fig. S2E). Discriminating morphology is therefore established early, well before the accuracy plateau, suggesting that full-length timelapses are not required for accurate classification (Fig. S2E). Extended trajectories nonetheless remain informative for resolving developmental dynamics, particularly in mutants exhibiting altered growth kinetics or biofilm dispersal defects. Together, these features distill each brightfield timelapse into interpretable feature trajectories that capture *V. cholerae* multicellular developmental and morphological phenotypes as accurately as the deep-learning embeddings.

### Genome-wide phenotyping reveals that emergent community morphology reflects cellular physiology

Given the utility of the µPULLI assay in discerning genotypes, we hypothesized that it could be applied more broadly to map genotype-to-phenotype relationships at scale. To this end, we screened an ordered *V. cholerae* transposon library of 2,850 non-essential genes, recording a 30-hour timelapse of every mutant (one image captured per hour, ∼88,350 images in total). Quantifying biofilm biomass trajectories revealed widespread phenotypic variation across the library: 53 strains (∼2%) exhibited reduced biofilm formation and 157 (∼6%) exhibited increased microcolony biomass or dispersal defects (Fig. 3A, Data S1, see Methods for thresholding strategy). As expected, many hits carried disruptions in canonical biofilm-associated pathways, including flagellar biosynthesis, polysaccharide matrix production, vibriobactin biosynthesis, and c-di-GMP signaling, confirming that the screen recovers known biological mechanisms underlying bacterial collective behaviors. Strikingly, however, the screen also uncovered distinct biofilm trajectories from the disruption of core physiological pathways not previously linked to biofilm regulation in *V. cholerae*, including tryptophan biosynthesis, zinc import, LPS/O-antigen biosynthesis, biotin biosynthesis, and pyruvate dehydrogenase flux (Fig. S4, Data S1). These results establish that our imaging assay can assign phenotypes across an entire non-essential gene set.

**Figure 3.**
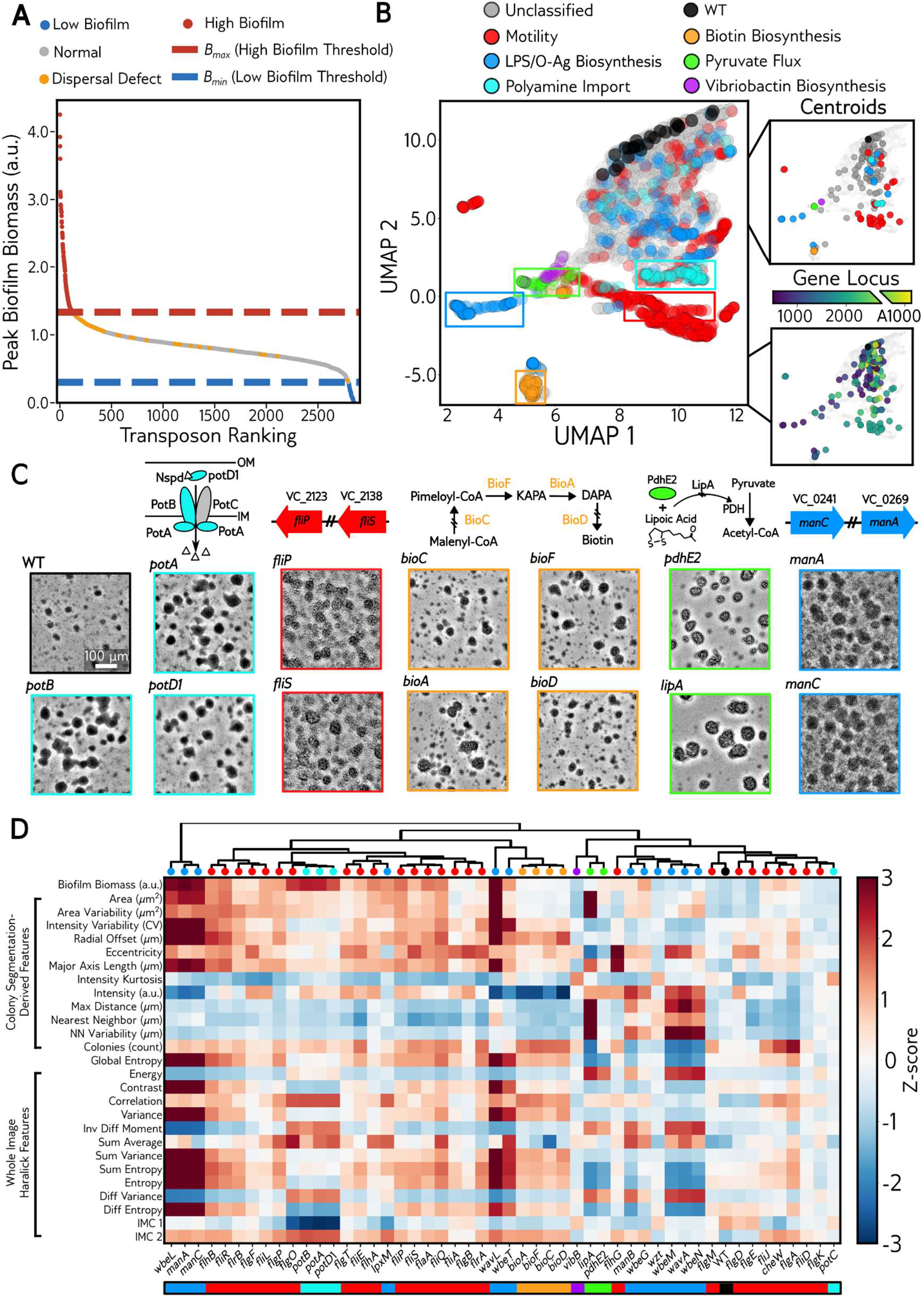
Genome-wide phenotyping with µPULLI-I reveals that emergent community morphology reflects cellular physiology. **(A)** Results from initial transposon screen. All mutants are ranked by normalized peak biofilm biomass and colored by phenotypic class (normal, low biofilm, high biofilm, or dispersal defect); dashed lines demarcate the High-Biofilm (red) and Low-Biofilm (blue) classification thresholds, with per-class counts in the legend. **(B)** UMAP dimensionality reduction of 157 *V. cholerae* High-biofilm/dispersal defect transposon insertion mutants (*N* ∼ 25 reimaging replicates/strain). Each point represents a single replicate. Plot is generated from biofilm biomass and whole-image quantitative features from 22 timepoints (9-30 h). Replicates are colored by their functional gene annotations. Rectangular boxes bound the replicates of the corresponding representative mutants shown in Fig. 3C, colored by functional annotation. Inset (top): Per-mutant centroids (mean UMAP coordinates across replicates), also colored by functional gene annotation. Each point represents a single transposon mutant from the 157 reimaged transposons. Inset (bottom): the same centroid positions as in the top inset, colored by chromosomal locus number (Chr. I, *VC_0001*–*VC_2756*; Chr. II, *VC_A0001*–*VC_A1099*) of transposon insertion. **(C)** Representative peak-biofilm brightfield images of mutants from each functional category, with pathway or operon schematics shown above each group; image border colors match the functional annotations in (B), and WT is shown for reference. Categories: polyamine import (*potA*, *potB*, *potD1*), motility (*fliP*, *fliS*), biotin biosynthesis (*bioC*, *bioF*, *bioA*, *bioD*), pyruvate flux (*pdhE2*, *lipA*), and O-antigen biosynthesis (*manA*, *manC*). Scale bar (white), 100 µm. **(D)** Hierarchical clustering dendrogram (top) and heatmap (bottom) of Z-scored biofilm features at each mutant’s respective peak biofilm biomass timepoint, for the subset of 50 transposon insertion mutants with displayed functional annotations (*N* = 25 replicates per mutant). Dendrogram (top) was computed by Ward linkage on PCA-reduced centroids (50 PCs) computed from the full feature-set trajectories for each mutant (see Methods). Circle markers on dendrogram leaves and colorbar below heatmap are colored by functional annotation. Feature values in the heatmap (bottom) are grouped by measurement type: colony segmentation-derived descriptors (colony count, nearest-neighbor distances), and Haralick textural statistics and Shannon entropy computed on the whole image. Color scale, Z-score relative to the full 157-mutant distribution. See Fig. S5A for complete dendrogram.

Many of the pathways above were hit by multiple independent insertions. Because mutants in the same functional pathways often converge on shared physiological defects at the single-cell level, we asked whether their phenotypes also converge at the level of multicellular community development and morphology, as captured by µPULLI-I. To assess this possibility, we generated up to 25 replicate videos per mutant of the 157 transposon hits with increased biofilm biomass or dispersal defects (3,867 videos after growth filtering, 119,877 images), and applied UMAP dimensionality reduction and hierarchical clustering to their extracted features, constructing a phenotype landscape of the biofilm-altered mutants (Fig. 3B–D). In UMAP space, many mutants formed a continuum around WT, while a subset resolved into discrete clusters comprising mutants with shared functional pathways (Fig. 3B). These groupings were also apparent by eye upon visual inspection of the images (Fig. 3C, Movie S2) and were recapitulated by hierarchical clustering performed directly on the extracted feature trajectories (Fig. 3D, Fig. S5A). The same groupings also emerged from UMAP dimensionality reduction and hierarchical clustering of DINOv2 embedding trajectories extracted with µPULLI-DL (Fig. S6A–C), indicating that the pathway structure is robust to how phenotypes are extracted and represented.

Across the dataset, genes within operons clustered closely together, including the polyamine import genes (*potA-D1*) and the biotin biosynthesis genes (*bioA-F*), while motility and LPS/O-antigen operons resolved into several sub-clusters (Fig. 3B, D, Fig. S5A). Notably, we observed that clustering was not restricted to co-located genes. For example, the pyruvate dehydrogenase mutant *pdhE2* co-clustered with *lipA*, which—despite being encoded in a distinct locus—supplies the lipoyl cofactor required by pyruvate dehydrogenase^31,32^ (Fig. 3C, D). Additionally, even in regions where the landscape appears continuous, structure persists to the level of individual genes, as replicates of the same genotype grouped far more tightly with each other than with replicates of other genotypes (Fig. S5B). Because our dendrogram (Fig. 3D) displays how each feature class varies across mutants at their peak biofilm biomass (though linkages are derived from full timelapse data), we also provide an accompanying animation that conveys how each feature evolves across the entire developmental time course (Movie S3, Interactive Plot 3). Moreover, interactive versions of the UMAP landscape and dendrogram link every replicate to its genotype and representative images (Interactive Plots 1-3). Together, these results show that genes converge in phenotypic space not only when co-located in an operon, but also when they contribute to the same physiological process from separate loci. Thus, community morphology reflects cellular physiology at a level deeper than gene co-location alone.

### Distinct phenotypes arise through transcriptional and non-transcriptional mechanisms

While our results suggest that cell physiology broadly influences collective behaviors and colony morphology, it is still unclear how disrupting different pathways produces distinct, pathway-specific phenotypes. Our transposon screen found that disruptions to biotin biosynthesis, pyruvate flux, and cell-surface architecture manifest as distinct phenotype clusters (Fig. 3, Fig. S6). Because these pathways are core to bacterial physiology and represent promising anti-infective targets,^33-36^ we selected a representative gene from each pathway for further characterization. To do so, we constructed clean, in-frame deletion mutants: Δ*bioD* (biotin biosynthesis; *VC_1115*), Δ*pdhE2* (pyruvate flux; *VC_2413*), and Δ*manA* (O-antigen biosynthesis; *VC_0269*). We then performed µPULLI-I for each strain and projected feature trajectories from in-frame deletion mutants onto the UMAP landscape constructed from feature trajectories from all 157 reimaged transposon mutants (Fig. 3B, 4A-B). Because the deletions are mapped onto the fixed transposon-reimaging UMAP embeddings rather than co-embedded, they do not reshape the landscape or move any transposon mutant. As expected, after projection each clean deletion mutant co-localized in UMAP space with its cognate transposon mutant (Fig. 4A), confirming that the phenotype flagged by the transposon screen reflects loss of the annotated gene rather than a transposon-specific or polar insertion effect.

**Figure 4.**
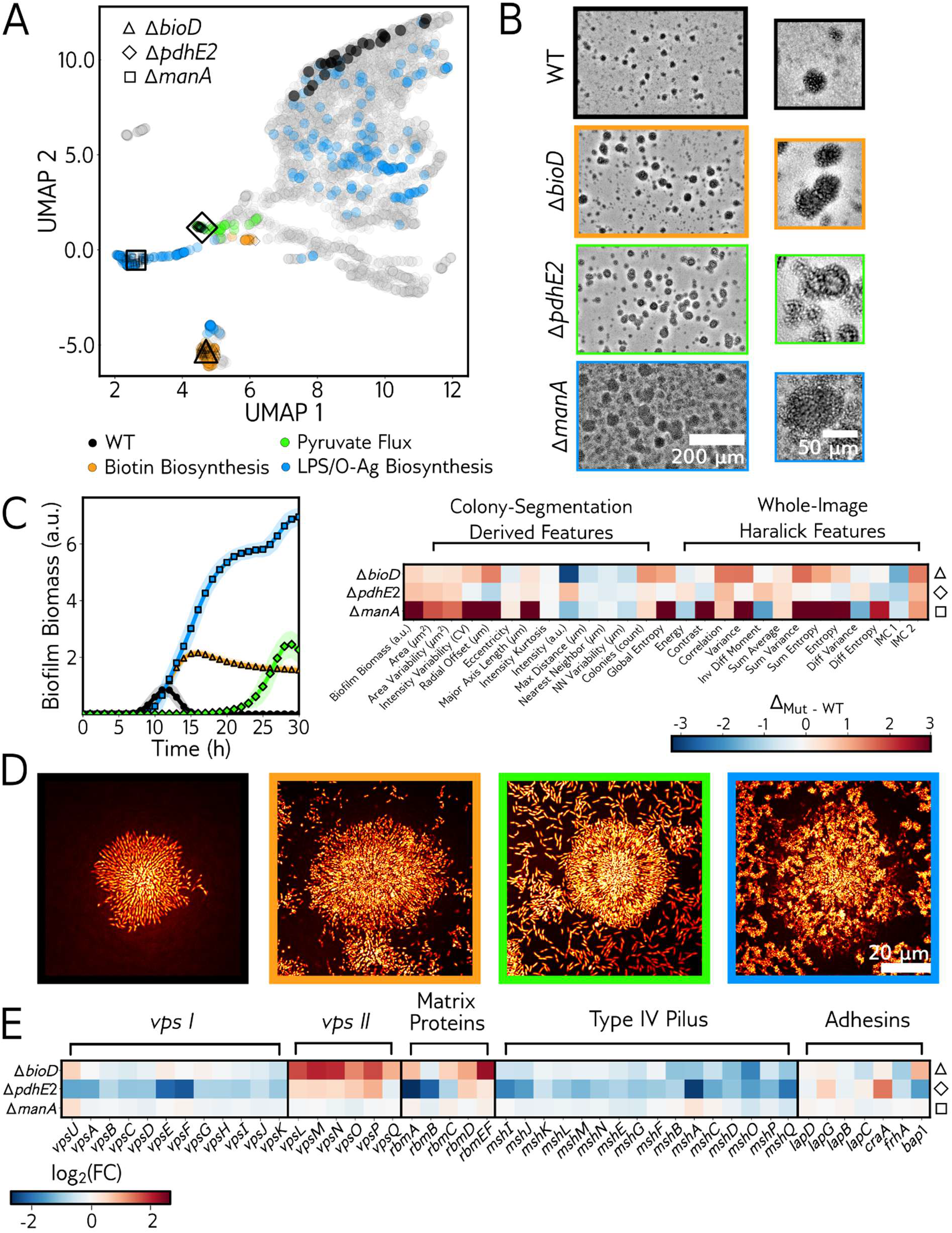
Metabolic and cell-envelope mutants drive distinct biofilm phenotypes through transcriptional and non-transcriptional mechanisms. **(A)** UMAP landscape projection of representative clean deletion mutants. Biofilm biomass and whole-image feature trajectories were extracted from 22 timepoints (9-30 h) of brightfield timelapse videos of clean, in-frame deletion mutants of representative genes from the biotin biosynthesis (Δ*bioD*, triangles), pyruvate flux (Δ*pdhE2*, diamonds), and LPS/O-antigen biosynthesis (Δ*manA*, squares) pathways. The clean deletion mutants were transformed into the fixed transposon reimaging embedding and projected onto the transposon reimaging UMAP landscape (Fig. 3B). Each gray point represents a single replicate from the transposon reimaging dataset (*N* = 24), and each open-face marker represents projected clean deletion mutants, as indicated (*N* = 3 biological, 3 technical replicates per clean deletion strain) where smaller open-face markers represent replicates, and each larger open-face marker represents a single centroid for all replicates of a strain. **(B)** Left: Representative brightfield images obtained at peak biofilm biomass for WT (12 h, black), Δ*bioD* (18 h, orange), Δ*pdhE2* (25 h, green), and Δ*manA* (18 h, blue). Scale bar, 200 µm. Right: Magnified inset from left panels. Scale bar, 50 µm. **(C)** Left: Biofilm biomass trajectories over 30 hours for WT (black), Δ*bioD* (orange), Δ*pdhE2* (green), and Δ*manA* (blue). *N* = 3 biological, 3 technical replicates. Shading represents ± s.d. Right: Heatmap of clean deletion mutant feature values at their corresponding peak biofilm-biomass timepoints relative to WT peak biofilm-biomass feature values. Each cell in the heatmap is colored by Δ_Mut - WT_ as a measure of scaled deviation from WT (see Methods). **(D)** High-resolution confocal fluorescence images of representative microcolonies (medial plane) for WT, Δ*bioD*, Δ*pdhE2*, and Δ*manA*. Cells were fixed at each mutant’s peak biofilm biomass timepoint (12 h, 18 h, 25 h, and 18 h, respectively), and stained with 500 µg/mL DAPI. Scale bar, 20 µm. See Fig. S7 for additional replicates. **(E)** RNA sequencing results displayed as heatmap focusing on biofilm-associated gene expression (log_2_ fold change relative to WT) for Δ*bioD*, Δ*pdhE2*, and Δ*manA*, grouped by functional category: *vps*-I, *vps*-II, matrix proteins, type IV pilus, and adhesins. See interactive plots 4-6 for display of full datasets. a.u., arbitrary units; FC, fold change.

Having confirmed that each representative in-frame deletion mutant reproduces its transposon phenotype, we examined what distinguishes the three mutants. Biofilm biomass trajectories revealed distinct developmental timelines for each mutant, differing in onset, peak biomass, and dispersal (Fig. 4C, left). Hallmarks of the Δ*bioD* phenotype include a ∼2-fold increase in peak biofilm biomass, reduced dispersal, and increases in colony number, size, size variability, and intensity. Δ*pdhE2* exhibited pronounced growth lag and a ∼2.5-fold increase in biofilm biomass, along with increased colony area and more uniform colony size. Part of the signature therefore reports altered growth kinetics, since the slowed growth rate delays the accumulation of Δ*pdhE2* biofilms (Fig. 4C). Meanwhile, Δ*manA*, the mutant with the most extreme biofilm phenotype, accumulated biofilm biomass steadily without appreciable dispersal, reaching ∼4-fold higher final biofilm biomass than WT (Fig. 4C). The Δ*manA* mutant was also highly enriched in colony number and size, and it showed increased variability of both colony size and intensity. Thus, deletions in genes in core physiological pathways at the cellular level cause distinct alterations to community-level morphology and collective behaviors, as detected by µPULLI-I.

To investigate whether phenotype differences identified by µPULLI-I are recapitulated at single-cell resolution, we imaged representative microcolonies using confocal microscopy (Fig. 4D). Each mutant’s brightfield signature corresponded to a distinctive biofilm architecture—densely packed biofilms for Δ*bioD*, a three-zone architecture for Δ*pdhE2* (a cell-dense core, a low-cell-density medial band, and a return of cell density toward the outer edge), and large fragmented biofilms surrounded by peripheral cell clumps for Δ*manA* (Fig. 4D, Fig. S7A). Thus, µPULLI can distinguish community-level phenotypic differences that are also apparent at single-cell resolution, without requiring labels or cell fixation.

Given the differences we observed in morphological and collective behaviors, we wondered if each phenotype arises from changes in biofilm gene expression or instead, whether phenotypic differences could arise from non-transcriptional changes to cell physiology. Because *V. cholerae* builds biofilms using numerous *Vibrio* polysaccharide (VPS) matrix genes (encompassing the *vps*-I and *vps*-II operons), matrix proteins, and adhesin genes, we reasoned that distinct biofilm gene expression programs could drive each unique phenotype.^37,38^ To examine this possibility, we performed RNA sequencing on the WT, Δ*bioD*, Δ*pdhE2*, and Δ*manA* strains (Fig. 4E, Fig. S7B, Interactive Plots 4–6). Indeed, Δ*bioD* and Δ*pdhE2* exhibited distinct changes in biofilm gene expression. For Δ*bioD* we observed strong upregulation of the *vps*-II operon and numerous matrix proteins compared to WT, consistent with its elevated biofilm biomass and dense biofilm architecture (Fig. 4D, E). The Δ*pdhE2* strain also exhibited a modest increase in *vps*-II expression, but surprisingly, displayed a significant decrease in the expression of *vps*-I genes and *rbmA*, which encodes a matrix protein that binds VPS and mediates cell-cell adhesion within microcolonies.^38,39^ Notably, however, *rbmB*—which encodes a polysaccharide lyase that trims the VPS matrix and is required for biofilm dispersal^40^—was downregulated ∼4-fold in the Δ*pdhE2* strain. We therefore propose that rbmB downregulation impairs matrix turnover, producing both the dispersal defect and the accumulation of biofilm biomass. In sharp contrast to the Δ*bioD* and Δ*pdhE2* strains, Δ*manA*, which showed the most divergent biofilm phenotype, exhibited no significant change in the expression of biofilm genes, and only 1.3% of genes were differentially regulated across the whole genome (Fig. 4E, Fig. S7B, Interactive Plots 4–6). Thus, the biofilm phenotype that deviates most from WT is invisible to transcriptomic assays. To validate our RNA sequencing results, we measured expression of a representative matrix gene (*vpsL*; first gene of the *vps*-II operon) using a luminescence reporter. Consistent with RNA sequencing, only Δ*bioD* and Δ*pdhE2* displayed significantly increased expression whereas Δ*manA* did not (Fig. S7C). These results demonstrate that distinct transcriptional changes underlie the Δ*bioD* and Δ*pdhE2* phenotypes, whereas the Δ*manA* phenotype cannot be explained by biofilm gene expression changes. We speculate that loss of O-antigen directly alters cell surface chemistry, driving attractive biophysical interactions between cells.^41^ Taken together, we conclude that genetic perturbations drive hallmark changes to biofilm development through both transcriptional and non-transcriptional mechanisms. Moreover, our results demonstrate that µPULLI captures biofilm states that are undetected by RNA sequencing.

### Emergent community morphology encodes rich, generalizable phenotypic information

Having established that community morphology reflects cellular physiology, we next asked whether µPULLI could characterize chemical perturbations agnostically by resolving the pathway a compound acts on from the phenotype it produces. Treating WT *V. cholerae* with saturating concentrations of MAC13772, an inhibitor of the biotin biosynthesis enzyme BioA,^42^ produced a phenotype indistinguishable from the Δ*bioD* deletion, and projecting the treated cultures onto the transposon-screen landscape placed them among the *bioA-F* biotin-biosynthesis transposon mutants (Fig. 5A, Movie S4). Reciprocally, supplementing Δ*bioD* with exogenous biotin restored a WT-like phenotype (Fig. 5A). These results indicate that µPULLI can be used to interrogate mechanisms underlying chemical perturbations, and in the future, could be used to nominate the target pathway of uncharacterized compounds by matching induced phenotypes against the phenotypes of genetic mutants.

**Figure 5.**
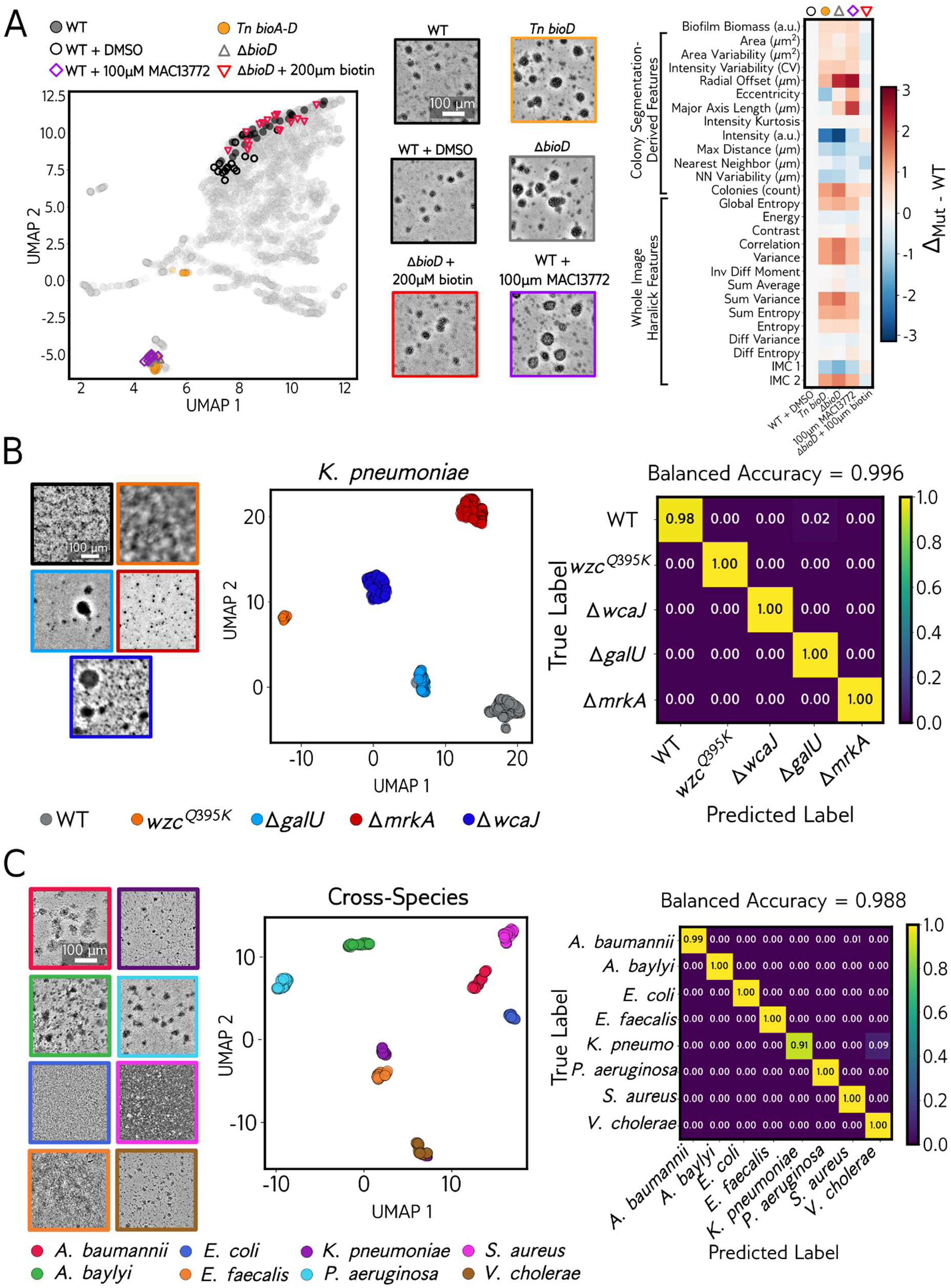
µPULLI resolves small-molecule mechanisms and generalizes across bacterial species. **(A)** Small-molecule modulation of biofilm development in *V. cholerae*. Left: UMAP phenotypic landscape (WT, dark gray; biotin-biosynthesis transposon mutants, orange) with overlaid conditions: WT + DMSO (open circles), WT + 100 µM MAC13772, a biotin biosynthesis inhibitor (purple diamonds), and Δ*bioD* + 200 µM biotin (red inverted triangles). *N* = 4 biological, 4 technical replicates per condition; 0.5% DMSO final concentration. Middle: representative brightfield images for each condition. Right: heatmap of biofilm features across conditions, expressed as Δ_Mut - WT_, quantifying normalized difference from WT (see Methods). **(B)** Extension to *K. pneumoniae*. Left: representative brightfield images captured using a 4× objective lens. Middle: UMAP of WT and four exopolysaccharide- and adhesin-associated mutants (*wzc^Q395K^*, Δ*wcaJ*, Δ*galU*, Δ*mrkA*). Right: confusion matrix for a random-forest classifier distinguishing the five strains from DINOv2 image embeddings, evaluated by repeated stratified 5-fold cross-validation over wells (5 repeats). Cultures were imaged for 24 hours; *N* = 96, 46, 96, 90, 96 wells for WT, *wzc^Q395K^*, Δ*wcaJ*, Δ*galU*, and Δ*mrkA,* respectively (424 wells total). Cells are row-normalized recall averaged over folds; mean balanced classification accuracy = 99.6%. **(C)** Species identification from monoculture growth. Left: representative brightfield images of the indicated species (*Acinetobacter baumannii*, *Acinetobacter baylyi*, *Escherichia coli*, *Enterococcus faecalis*, *Klebsiella pneumoniae*, *Pseudomonas aeruginosa*, *Staphylococcus aureus*, and *Vibrio cholerae*) at their peak biofilm biomass timepoints, captured using a 10× objective lens. Middle: UMAP dimensionality reduction of DINOv2 embeddings generated from 16 timepoints (9-24 h) of brightfield timelapses of the eight bacterial species. Right: confusion matrix for species classification from DINOv2 image embeddings using a random forest classifier, evaluated by plate-grouped 5-fold cross validation (5 repeats). For consistency, all organisms were grown in LB medium at 30 °C and brightfield timelapses were captured for 24 hours. *N* = 18–30 wells per species (210 total). Cells are row-normalized recall averaged over folds; mean balanced accuracy = 98.8%.

We next asked whether the µPULLI approach generalizes beyond *V. cholerae*, specifically, using the deep-learning approach (µPULLI-DL) as it requires no segmentation and can be used to extract phenotypes from images of any organism. Applied to the biofilm-forming pathogen *Klebsiella pneumoniae*,^43-45^ µPULLI-DL distinguished WT and four exopolysaccharide- and adhesin-defective mutants with near-perfect accuracy (cross-validated balanced classification accuracy 99.6%; Fig. 5B, Movie S5), showing that the approach can be extended without modification to dissect the phenotypic impact of genetic perturbations in other pathogens. Scaling further, we asked whether µPULLI-DL could assign species identity from low-magnification videos of growing monocultures. Indeed, µPULLI-DL separated eight taxonomically diverse species, spanning Gram-negative and Gram-positive organisms, with near-perfect accuracy (cross-validated balanced classification accuracy = 98.8%; Fig. 5C, Movie S6). Together, these results demonstrate that the emergent morphology of a bacterial culture, imaged at low magnification and without labels, carries sufficient information to identify the pathway targeted by chemical inhibitors, resolve genetic mutants in other pathogens, and distinguish diverse species. Reading this collective-scale structure thus provides a general route to phenotyping bacteria from simple, low-magnification brightfield images.

## DISCUSSION

Bacterial phenotyping has long relied on assays that are committed in advance to what they measure. What such assays can discover is therefore bounded by the phenotypic traits they are optimized to detect, and phenotypes falling outside that expectation go unrecorded. This constraint has become more limiting as isolates and mutants can be sequenced faster than they can be functionally characterized.^7,46^ Image-based profiling avoids this constraint by recording morphology first and determining afterward which properties are informative.^47^ The approach has become a mainstay in mammalian cell biology via Cell Painting^10^, where fluorescent stains resolve organelles and other subcellular structures and perturbations manifest as a change in a cell’s internal organization. Even label-free implementations, which predict Cell Painting channels from brightfield images,^48^ are capable of reading out subcellular organization. Bacteria present no comparable interior organization at conventional resolution and consequently, label-free bacterial imaging has primarily remained focused on cross-species classification and shape defects at the scale of the single cell.^49-53^ What bacteria offer instead is immense phenotypic richness and diversity in the emergent multicellular architectures that arise from population-level organization.

Humans have long relied on emergent form as a carrier of identity. Tasks such as recognizing individual faces and distinguishing plant and animal species based on visual inspection alone are now also performed by learned image models.^54-56^ Principally, such approaches are effective because a multicellular organism’s developmental trajectory and overall form are heavily influenced by its genotype. Our work demonstrates that, similarly, microbial multicellular collectives carry significant phenotypic, and therefore genotypic, information, perhaps more than can be gleaned from examination of single cells. Bacterial colony morphology has long served this purpose on agar petri plates, where rugose and smooth variants are distinguished by eye and learned models now assign strain identity from colony images.^57,58^ Colonies on agar, however, are historically scored categorically and generally at a single endpoint. Growth in liquid culture, by contrast, allows for the continuous examination of microcolony growth and dispersal dynamics. Moreover, such cultures can be seeded by simple liquid handling protocols, and cultures in microwell plates are readily parallelized for imaging, enabling large-scale phenotyping efforts. Our imaging approach, <u>Micro</u>bial <u>P</u>henotyping <u>U</u>sing <u>L</u>ow-magnification <u>L</u>abel-free Imaging (µPULLI), defines emergent community architecture as a continuous, time-resolved phenotype by simply imaging bacterial growth in liquid cultures at low-magnification as they grow over time.

Using µPULLI, we measured multicellular form agnostically and demonstrated that genotype influences collective behavior and morphology more thoroughly than any single standard biofilm assay would suggest. Biofilm biomass, the single property measured by conventional crystal violet assays,^8^ was sufficient on its own to distinguish eight genetically distinct *V. cholerae* strains at 93.8% balanced classification accuracy when resolved as a dynamic developmental trajectory rather than an endpoint measurement. Colony morphology and image texture were each also independently sufficient to discriminate genotypes, and crafting a full feature set comprising biofilm biomass, colony morphology, and whole-image texture statistics together achieved near-perfect classification accuracy on the 8-strain *V. cholerae* panel. We also demonstrated that genotype identity is not restricted to one visual property or feature class, but distributed redundantly across the amount of biofilm a culture forms as well as the shape, arrangement, and texture of its microcolonies. Because genotypic information is encoded independently across feature classes, and because biofilm biomass and texture are measured without resolving individual microcolonies, genotype remains recoverable when segmentation fails. The same redundancy explains why µPULLI-DL was equally effective at discriminating strains: a general-purpose encoder trained on natural photographs, with no segmentation and no fine-tuning, matched the accuracy of features we designed by hand. Thus, genotypic signal is distributed broadly enough in low-magnification brightfield images of multicellular communities that it does not depend on measuring any particular property.

Applied across the *V. cholerae* transposon library, µPULLI revealed that roughly 8% of genes altered how the population assembles. Importantly, many of our hits had no previously established connection to biofilm formation in this organism. Disruptions to genes in biotin synthesis, pyruvate flux, and O-antigen production pathways each manifested in reproducible, distinctive multicellular phenotypes, demonstrating that bacterial multicellular community development and morphology are greatly influenced by cellular physiology. Biofilm development and morphology can therefore be thought of as an integrator of cellular state—unsurprising in one respect, since transcription factors connect cellular states to biofilm matrix production through broad transcriptomic alterations.^37,59^ However, the Δ*manA* strain shows the integration of cellular physiology in community architecture is not limited to a transcriptional route. We demonstrated that the Δ*manA* mutant produced the most divergent biofilm phenotype, yet left essentially no transcriptional signature; a discrepancy we hypothesize is due to the biophysical consequences of losing O-antigen rather than to altered gene expression.^41^ Thus, µPULLI revealed that multicellular phenotypes can be extreme without involving gene regulation, and the extent to which bacterial physiology is legible in community form while invisible to transcription remains to be discovered.

Bacteria have been grown in liquid culture and observed under microscopes for more than a century, and for most of that time the developmental information contained in those cultures has been underutilized. µPULLI closes that gap and provides a route for rapid phenotyping in the genomic era. Because we discovered—using unsupervised methods—that mutants grouped phenotypically by the pathways they disrupted, our datasets support direct inference: an uncharacterized perturbation can be interpreted from the known mutants it most resembles.^60,61^ We also demonstrated that the operation is not specific to genetic mutants of *V. cholerae*. Projecting into the same coordinate system of genetic perturbations assigned a small molecule to its target pathway from images alone, separated *K. pneumoniae* mutants, and resolved eight taxonomically diverse species, none of which required modification to the platform. Reading collective form therefore requires no reporters, no genetic tractability, and no prior model of how an organism grows, which is what makes a single reference set extensible rather than organism-bound. As such, µPULLI can be applied to clinical isolates, laboratory evolution experiments, and mixed communities. Imaging additional growth conditions would thereby extend the same reference set rather than create separate ones. We release the atlas, interactive viewers, analysis code, and liquid-handling protocols so that others can add to it.

### Limitations of the study

The V. cholerae landscape was constructed under a single growth condition, in one medium at one incubation temperature, so we do not know how it varies with the environment. Most phenotypes will likely shift to some degree under different conditions, and which groupings persist can only be established by re-measuring the panel in each new condition. Coverage of the map is also incomplete: the landscape was built from replicate imaging of the 157 mutants with increased biofilm biomass, delayed biofilm formation, or dispersal defects; essential genes lie outside the transposon library entirely, and insertional mutagenesis carries the usual risk of polar effects on neighboring genes. Co-clustering indicates a shared physiological state rather than a shared mechanism, and while the *pdhE2* and *lipA* association recovered by µPULLI is corroborated by known biochemistry,^31,32^ most associations in the map are hypothesis-generating and require independent validation. The same caution applies to *manA*, where we infer that the altered multicellular community development and morphological phenotype arose from the accompanying loss of O-antigen and the peripheral clumping we observed, although the biophysical mechanism itself has not been demonstrated. Finally, UMAP serves as a visualization tool only in this study; the UMAP embeddings do not inherently preserve distance, and proximity in a projection is not evidence of phenotypic similarity. Relatedness claims here rest on hierarchical clustering and permutation tests computed in the original feature space.

## METHODS

### Bacterial Strains and Growth Conditions

The parent *V. cholerae* strain used in this study was WT O1 El Tor biotype C6706str2. The *V. cholerae* ordered transposon mutant library was a gift from the laboratory of Chris Waters at Michigan State University.^17^ *E. coli* S17 and Top10 were used to transfer plasmids into *V. cholerae* by conjugation. For propagation and cloning, strains were grown in lysogeny broth (LB) with shaking (200 RPM), or on LB plates supplemented with 1.5% agar, at 30 °C. For microscopy, luminescence quantification, and RNA sequencing, *V. cholerae* cells were grown in M9 medium containing glucose and casamino acids (1× M9 salts, 100 µM CaCl_2_, 2 mM MgSO_4_, 0.5% glucose, and 0.5% casamino acids). For imaging of the *K. pneumoniae* five-strain panel (Fig. 5B), strains in the KPPR1 background were grown overnight with shaking in LB and subsequently back-diluted into M9 medium containing glucose (1× M9 salts, 100 µM CaCl_2_, 1 mM MgSO_4_, 0.4% glucose) for microscopy of static growth. As described previously, this *K. pneumoniae* M9 medium was chelated with Chelex-100 resin for 3 hours to remove trace metals, and salts were re-added after filter sterilization.^29,62^ For the multispecies panel (Fig. 5C), all other species were grown overnight and imaged in LB. For all *V. cholerae* and *K. pneumoniae* experiments, overnight cultures were grown at 37 °C, and imaging was performed at 30 °C. All other species were pre-cultured and imaged at 30 °C. Unless otherwise noted, antibiotics were used at the following concentrations: kanamycin, 50 µg/mL; spectinomycin, 200 µg/mL; streptomycin, 400 µg/mL; chloramphenicol, 2 µg/mL; ampicillin, 100 µg/mL. For treatment with biotin (Fisher Scientific) and MAC13772 (Med Chem Express), compounds were dissolved in DMSO and added at a final concentration of 200 µM and 100 µM, respectively, to the indicated strains at the time of inoculation. All strains used in this study are reported in Table S2.

### Genetic Manipulation and Strain Construction

All strains constructed in this study were generated by replacing genomic DNA with linear DNA introduced by natural transformation, as described previously.^63,64^ Linear DNA fragments were generated by splicing overlap extension (SOE) PCR using iProof (Bio-Rad) or Q5 (New England Biolabs) DNA polymerase, joining products amplified from *V. cholerae* genomic DNA. In all cases, approximately 3 kb of upstream and downstream flanking homology was included to promote efficient chromosomal integration. Oligos were ordered from IDT and are reported in Table S3.

To induce natural competence, the parent strain was grown overnight from a single colony at 30°C in liquid LB with agitation, diluted 1:1,000 into fresh LB, and grown to an OD_600_ of ∼1.0. Cells were pelleted and resuspended in an equal volume of 1× Instant Ocean sea salts (7 g/L). A 100 µL aliquot was added to 900 µL of a chitin–Instant Ocean suspension (8 g/L chitin, Alfa Aesar) and incubated overnight without agitation at 30 °C. The next day, the linear DNA fragment carrying the desired alteration was added together with a selectable antibiotic-resistance cassette co-integrated at the neutral locus *VC_1807*, and the mixture was incubated for 12–24 hours at 30°C. Cells were recovered in liquid LB for one hour, plated on LB agar containing the relevant antibiotics, and restreaked on selective LB plates. Clean, in-frame deletions of *bioD*, *pdhE2*, and *manA* were generated to remove the entire coding sequence, and all modifications were verified by PCR and Sanger sequencing (Azenta). The P*_vpsL_-lux* reporter plasmid was introduced separately by triparental mating with *E. coli* S17 and Top10.

### Inoculation and Liquid Handling Protocols for µPULLI

To ensure reproducibility across all µPULLI assays, imaging plates were seeded on an Opentrons OT-2 liquid handling robot using custom protocols. In previous work, we found that seeding at low starting cell densities (i.e., 10^4^–10^5^ CFU/mL) enabled robust microcolony biofilm formation across diverse species.^25^ Seeding therefore requires a large dilution from the stationary-phase overnight culture, which we achieved by serial dilution through a 384-well intermediate plate. Strains to be imaged were first grown overnight in 200 µL of LB in polystyrene 96-well plates (Corning #3370) with shaking at 37 °C for 24 hours, covered with a Breathe-Easier membrane (USA Scientific). The following day, to ensure accurate liquid handling, pellicle material at the air-liquid interface was removed using the bottom of a sterile 96-well PCR plate, whose conical points adhere to surface pellicle, and any residual film was cleared with a pipette tip. To perform the dilution series, the OT-2 deck was loaded with the overnight-culture plate, 20 µL and 300 µL pipette tips (Opentrons), a 12-well media reservoir (USA Scientific), a sterile 384-well plate (NEST Scientific), and destination 96-well plates (same polystyrene model as above) to be used for microscopy. In the liquid handling protocols, each overnight culture was serially diluted 1:20 in imaging media (M9 or LB) four times in the 384-well plate (50 µL per well) without changing pipette tips. After a tip exchange, a final 1:60 dilution was performed to inoculate the 96-well imaging plates in 150 µL of imaging media per well. At each transfer step, cultures were mixed on the OT-2 by five aspirate–dispense cycles of 20 µL. Although the calculated dilution factor was high, we experimentally determined that the final CFUs were comparable to the cell dilutions previously used in our lab,^25^ presumably due to cell retention on the pipette tips. For the initial transposon screen, each overnight plate harboring unique transposon mutants was used to seed a single imaging plate. For all replicate imaging (*V. cholerae* known mutants, Fig. 1–2; transposon re-imaging, Fig. 3; *K. pneumoniae* and other species, Fig. 5), a single overnight 96-well plate containing biological strain replicates was used to inoculate three 96-well technical replicate imaging plates in a single OT-2 run. For *K. pneumoniae*, the first two dilutions in the 384-well plate were instead performed in PBS, and the second two in M9. All Opentrons protocols are available on our Kilthub repository.

### Brightfield Microscopy and Robotic Plate Transfer

Brightfield timelapse images were acquired on an Agilent BioTek Cytation 5 imaging plate reader driven by Gen5 software (version 3.12.08). Inoculated 96-well plates were incubated in an attached Agilent BioTek BioSpa 8 automated incubator (BioSpa OnDemand software) at 30 °C, and the BioSpa robotic arm transferred each plate to the Cytation 5 for imaging every hour. The *V. cholerae* datasets were imaged over a 30-hour period, yielding 31 frames per well; the *K. pneumoniae* and multispecies datasets were imaged over 24 hours. Images were captured with a 10× air objective (Olympus Plan Fluorite, NA 0.3), except for *K. pneumoniae*, which was imaged at 4× (Olympus Plan Fluorite, NA 0.13). Focal position was maintained with laser autofocus. Exposure, illumination intensity, and gain settings were held constant across all compared datasets.

### Quantification of P*_vpsL_-lux*

To monitor *vpsL* expression as a proxy for biofilm matrix production (Fig. S7C), we leveraged a P*_vpsL_-lux* transcriptional reporter, in which the *vpsL* promoter is fused to the *luxCDABE* luciferase operon of *Photorhabdus*.^65^ Three biological replicate overnight cultures were back-diluted 1:5,000 in fresh M9 medium (OD_600_ of ∼10^-4^) and seeded in triplicate in 96-well plates (Corning). Optical density (OD_600_) and luminescence were then recorded in parallel at 1-hour intervals over 24 hours on an Agilent BioTek Cytation 5 plate reader running Gen5 software at 30 °C.

### Spinning Disk Confocal Microscopy

For confocal imaging, cells were grown in glass-bottom 96-well plates (Mattek) in M9 media to the timepoint at which microcolony biomass peaks (12 h for WT, 18 h for Δ*bioD* and Δ*manA*, and 25 h for Δ*pdhE2*). Biofilms were then fixed in 3.7% formaldehyde (in 1× PBS) for 20 min, washed three times in 1× PBS, and stained with 500 µg/mL 4′,6-diamidino-2-phenylindole (DAPI) for 1 hour before imaging. Images were acquired on a motorized Nikon Ti-2E microscope fitted with a CREST X-Light V3 spinning disk unit and a back-thinned sCMOS camera (Hamamatsu Orca Fusion BT), using a 100× silicone immersion objective (Nikon Plan Apochromat, NA 1.35). DAPI was excited at 405 nm, acquisition was controlled by Nikon Elements software (version 5.42.02), and illumination was provided by an LDI-7 Laser Diode Illuminator (89-North).

### RNA Isolation and Transcriptomic Analysis

For transcriptomic analysis, three biological replicates of each strain were grown overnight, back-diluted 1:5,000 into 20 mL M9, shaken at 30 °C, and harvested at OD_600_ = 0.1. Cells were collected by centrifugation (3,200 × g, 10 min), and total RNA was isolated with the Monarch Total RNA Miniprep Kit (New England Biolabs), including on-column DNase treatment to remove genomic DNA. RNA concentration and purity were measured on a NanoDrop instrument (Thermo). Samples were flash-frozen in liquid nitrogen, stored at −80 °C, and shipped on dry ice to SeqCenter (https://www.seqcenter.com/rna-sequencing/). Following rRNA depletion, samples were sequenced to 12 M paired-end reads with the intermediate analysis package. As reported by SeqCenter, quality control and adapter trimming were performed with bcl-convert (Illumina; 4.2.4; default parameters), read mapping with HISAT2 (2.2.0; default parameters + ‘--very-sensitive’),^66^ and read quantification with Subread’s featureCounts (2.0.1; default parameters + ‘-Q 20’).^67^ Counts were loaded into R (R Core Team; 4.0.2; default parameters) and normalized by edgeR’s (1.14.5; default parameters)^68^ trimmed mean of M-values (TMM) method, then converted to counts per million (CPM). Differential expression was assessed with edgeR’s glmQLFTest, and genes with |log_2_ fold-change| > 2 and *P* < 0.05 (Benjamini-Hochberg) were considered significant. Heatmaps were generated with Seaborn and Matplotlib in Python. Interactive volcano plots were generated with a custom script combining native JavaScript and the HTML5 Canvas API, with data processing and log-transformation in Python (Pandas and NumPy).

### Brightfield Image Preprocessing

Brightfield timelapse images were preprocessed using a custom Python pipeline (github.com/BridgesLabCMU/uPULLI-I) that extends our previously described method written in Julia for automated multi-well biofilm imaging, label-free analysis of biofilms (LFAB).^29^ For each frame, raw pixel values were scaled to [0, 1] according to the acquisition bit depth (16-bit). To suppress large-scale illumination gradients while preserving local biofilm contrast, each frame was high-pass normalized: a local background was estimated as a Gaussian low-pass (σ ≈ 101 px, computed on an 8×-downsampled image for efficiency), and the frame was subtracted from this background so that dark biofilm renders as positive contrast. The resulting residual was lightly smoothed with a Gaussian filter (σ = 2 px). Frame-to-frame drift was corrected by Fourier-domain phase correlation (OpenCV phaseCorrelate), estimating the sub-pixel translational offset from the location of the cross-power-spectrum peak; a shift was computed for every frame relative to the preceding frame, the per-frame shifts were accumulated, and any proposed shift exceeding 250 px was discarded as an acquisition artifact. The accumulated shifts were applied by affine translation, and border rows and columns vacated by registration were removed by cropping.

### µPULLI-I Microcolony Biofilm Segmentation and Biomass Quantification

Binary biofilm masks were generated by thresholding the processed images at an empirically determined linear intensity threshold of 0.025 for all *V. cholerae* datasets in this manuscript (Figs. 1–4), 0.04 for the *K. pneumoniae* known-mutant dataset imaged using a 4× air objective lens (Fig. 5B), and 0.03 for the multispecies classification dataset imaged using a 10× air objective lens (Fig. 5C). Dust correction was then applied, such that persistent bright artifacts—defined as pixels present in the mask at frame 0 but absent in at least one subsequent frame—were removed across all frames. Per-frame biofilm biomass was computed from optical density referenced to each well’s own biofilm-free baseline. For each well, a per-pixel blank I_blank was taken as the mean of the first five frames (which precede the formation and appearance of biofilms), and optical density was computed as OD_t(x,y) = −log₁₀(I_t(x,y) / I_blank(x,y)) on the registered raw intensities. Biomass at each frame was the sum of OD over masked pixels, normalized by total image area. Referencing each well to its own blank frames cancels per-well and per-session differences in exposure and illumination, while the measure continues to scale with both biofilm coverage and optical contrast relative to the clear-well background.^29^

### Individual Microcolony Biofilm Tracking and Segmentation

To obtain features of individual biofilms, microcolonies were tracked over time by label propagation from a seed frame, defined as the point at which microcolonies first become visible in the processed, locally normalized images. A nearest-neighbor label propagation approach was developed specifically for this dataset because of the nature of *V. cholerae* biofilms: surface-attached biofilms expand radially and frequently contact and fuse at their boundaries. Because these fused structures share no resolvable intensity boundaries, traditional segmentation methods such as watershed or contour-based algorithms consistently oversegment or incorrectly merge colonies at fusion events. Distance-transform-based propagation instead assigns boundary pixels to the nearest existing labeled region rather than requiring an intensity edge, allowing colony identities to be maintained through fusion events by spatial proximity alone.

The seed frame was defined as the first frame at which per-frame biofilm biomass exceeded 0.005 for at least two consecutive frames, while the peak frame was defined as the frame of maximum computed biofilm biomass. Prior to labeling, each frame’s binary colony mask was hole-filled (SciPy ndimage binary_fill_holes, v1.13.1) and connected components smaller than 200 pixels were removed. From the seed frame onward, each pixel in the binary mask was assigned the label of its nearest previously-labeled pixel using a distance transform, with an effective propagation radius that scaled with the inter-frame time gap before the peak frame and was held fixed thereafter (at 50 pixels for the *V. cholerae* datasets which were imaged with the 10× objective). Pixels not claimed by an existing label were grouped into connected components: components meeting the 200-pixel minimum area were assigned new colony identities and smaller fragments were discarded, preventing mask-noise fragments from acquiring spurious identities.

To keep colony identities stable when growth stalls, labels were propagated against a persistent label footprint rather than the immediately preceding frame alone. When colonies are not actively expanding, binary mask edges flicker due to thresholding noise, so pixels that transiently drop below threshold and later reappear can fall outside the propagation radius of the previous frame and acquire spurious new identities under naïve propagation. To prevent this, the propagation reference was a cumulative footprint that never shrinks: after each frame, newly labeled pixels were added to the footprint and existing labels were never overwritten, so a recovered pixel was re-attracted by the distance transform to the label it had previously held. Tracking quality was confirmed by visual inspection across a representative subset of wells. Colonies and connected components along the image border were excluded from downstream analysis.

### µPULLI-I Feature Extraction

Two classes of features were extracted per well per frame: whole-image features and colony-level features.

Whole-image features were computed from the processed image stacks and included the following pixel-intensity statistics (Table S1): Shannon entropy and the 13 Haralick co-occurrence texture features^30^ that describe textural statistics over the gray-level co-occurrence matrices for each frame (mahotas, v1.4.18).

Colony-level features comprised individual colony geometric properties, intensity statistics, and inter-colony relationships, extracted per colony per frame (Table S1). Colony identity and geometry were taken from the tracked label stack, while intensity features were measured from the processed image stack (the fixed-background render also used for the whole-image features), not the raw images. Geometric properties included colony area, radial offset, and eccentricity; all colony-level geometric quantities are expressed in µm or µm^2^ using a conversion factor of 0.697 µm pixel^-1^ for the 10× objective hardware. Intensity properties included mean, integrated, and maximum intensity, as well as the offset between the geometric and intensity-weighted centroids. Inter-colony spatial properties included nearest-neighbor distances (k = 1, 5) and minimum spanning tree edge statistics across colonies in the field of view.

Colony features were aggregated to the well level as the mean, standard deviation, skewness, and kurtosis across all colonies per frame. Per-well biomass time series and feature tables were assembled into plate- and experiment-level CSV files for downstream analysis.

### µPULLI-DL Vision Transformer Embeddings

All frames of the preprocessed brightfield timelapse stacks (local-contrast normalized, dust corrected, registered, and cropped to 1,992 × 1,992 pixels) were used as input to a pretrained vision transformer (DINOv2 ViT-B/14). Frames were read from the processed image render, which is scaled to [0, 1], and each frame was resized to 518 × 518 pixels by bicubic interpolation (the native input resolution for DINOv2 ViT-B/14). Single-channel grayscale frames were replicated across three channels and standardized with the ImageNet channel statistics used in DINOv2 pretraining (mean [0.485, 0.456, 0.406], standard deviation [0.229, 0.224, 0.225]) to match the model’s expected 3-channel input distribution.

Frame-level features were extracted using DINOv2-Base (ViT-B/14), a vision transformer pretrained with self-supervised DINO objectives on ∼142 million images.^27^ The model was used as a frozen feature extractor without fine-tuning. At 518 × 518 pixel input resolution, the patch embedding layer divides each frame into a 37 × 37 grid of non-overlapping 14 × 14 pixel patches, each projected to a 768-dimensional token. From the final transformer layer we extracted two complementary representations per frame: (i) the classification (CLS) token, a single 768-dimensional vector summarizing the global visual state of the well, and (ii) the patch tokens, which encode spatially localized regions and were pooled from the 37 × 37 grid to a 3 × 3 grid by adaptive average pooling, yielding nine 768-dimensional regional descriptors per frame.

Each well was represented as the temporal trajectory of its per-frame CLS-token vectors—one 768-dimensional vector per timepoint—concatenated in order into a single fixed-length vector, preserving the full temporal trajectory of the well’s global visual state without collapsing its temporal structure. Trajectory length was fixed per organism: *V. cholerae* wells used a common 31-frame trajectory (trajectories longer than 31 frames were truncated to their first 31 so that same-organism trajectories were length-matched for comparison and UMAP projection), while *K. pneumoniae* defined-mutant wells (Fig. 5B) used their full 25-frame trajectories, and the multispecies classification wells (Fig. 5C, 10×) used 25-frame trajectories. The concatenated representation was therefore 31 × 768 = 23,808 dimensions for *V. cholerae*, 25 × 768 = 19,200 dimensions for *K. pneumoniae*, and 25 × 768 = 19,200 dimensions for the multispecies dataset. Because the trajectory length differs, these embedding sets are not dimension-compatible and were analyzed as separate representation spaces.

### Dataset Filtering

Replicates from the initial *V. cholerae* eight-strain dataset comprising (Fig. 1 and 2) of WT, Δ*vpsL*, Δ*rbmB*, Δ*hapR*, Δ*potD1*, Δ*flaA*, *luxO^D61E^*, and *vpvC^W240R^*, the 157-mutant transposon reimaging set (Fig. 3, 4), and the five-strain *K. pneumoniae* mutant dataset (Fig. 5) comprising WT, *wzc^Q395K^*, Δ*wcaJ*, Δ*galU*, and Δ*mrkA*, were subjected to a growth filter to exclude any replicates that did not grow or produce biofilm biomass. Specifically, a replicate from any dataset was excluded if it did not produce biofilm biomass ≥ 15% of the median peak biomass of the WT replicates on the same plate (or the dataset’s WT plate where WT was imaged separately). The non-biofilm forming Δ*vpsL* was excluded from this filter in all *V. cholerae* datasets.

### Dimensionality Reduction and Visualization

Dimensionality reduction was performed with UMAP^28^ (umap-learn v0.5.9), treating each timelapse replicate as a single observation and coloring points by strain, species, or functional annotation as indicated. Two feature representations were used in parallel: (i) For µPULLI-I, hand-engineered image features (biofilm biomass, whole-image Haralick and Shannon-entropy features, and—for the defined mutant set (Fig. 2)—colony-segmentation-derived features), and (ii) For µPULLI-DL, DINOv2 CLS embeddings, formed by stacking the per-frame CLS token over the growth-phase window into a single vector per well. Feature matrices and the Euclidean embedding were standardized to zero mean and unit variance (StandardScaler, scikit-learn v1.6.1). Replicates were growth-filtered identically to the classification analyses (see *Phenotypic classification and cross-validation*). The single-dataset UMAPs (Fig. 1D, 2D, 5B, 5C) were each fit *de novo* on their dataset. Hyperparameters and timepoint windows for all datasets are given in the corresponding figure legends.

*Reimaging manifold and projection:* The *V. cholerae* transposon reimaging dataset manifold—the shared coordinate space onto which the compound-treatment and clean-deletion datasets are projected (Fig. 3, 4, 5A)—was built once from biofilm-biomass and whole-image (Haralick and Shannon-entropy) features at timepoints t=9–30 hours (colony segmentation-derived features excluded), standardized as above. UMAP embeddings were computed over a 3 × 3 grid of n_neighbors ∈ {10, 20, 30} and min_dist ∈ {0.1, 0.2, 0.3}, with n_components = 2, metric = ‘euclidean’, and random_state = 0. The canonical landscape in Fig. 3 used n_neighbors = 10, min_dist = 0.1. The fitted StandardScaler, UMAP model, and feature-column list were saved so that the clean-deletion and compound datasets (Fig. 4, 5A) could be projected into that same space via UMAP.transform() without refitting. Before projection, projected datasets were subjected to a growth filter and were transformed directly without batch correction. Projection fidelity was quantified by the Euclidean distance between each projected clean-deletion mutant’s centroid and its transposon counterpart’s centroid in the manifold.

*Interactive UMAP:* To let readers explore the manifold, we provide an interactive UMAP as a self-contained supplemental file (Interactive Plot 1): a single portable HTML document with all imagery embedded, so it runs offline without a server. Users can switch among the full n_neighbors × min_dist parameter grid to see how these hyperparameters reshape the manifold, toggle functional-annotation highlights, search for individual genes, and pan/zoom; clicking any point displays that replicate’s peak biofilm biomass image (the maximum-biomass frame of its processed timelapse).

### Phenotypic Classification and Cross-Validation

To assess how well µPULLI distinguishes the eight *V. cholerae* defined mutant genotypes (see Fig. 1E, 2E), we trained random-forest classifiers (200 trees, min_samples_leaf=2, class_weight=‘balanced’, random_state=0; scikit-learn v1.6.1) under a common cross-validation protocol and applied them to two feature representations. (i) Quantitative imaging features: biofilm biomass, whole-image entropy and Haralick features, and colony segmentation-derived features, evaluated both as the full feature set and as each of seven feature-class combinations. (ii) DINOv2 CLS embeddings: the per-frame CLS token was stacked over the growth-phase window (timepoints 9–30 hours) into a single vector per well. In both cases, performance was estimated with GroupKFold cross-validation using imaging plate as the grouping variable (5 folds × 5 repeats, with plate-fold assignments reshuffled between repeats), so that all replicate wells from a plate were held out together and accuracy reflects generalization across plates rather than plate-specific effects. Features were standardized (StandardScaler) using statistics fit on the training fold only. Performance was summarized as balanced classification accuracy (mean per-class recall, robust to unequal replicate counts across genotypes); confusion matrices were row-normalized within each fold and averaged across all folds and repeats. Wells were retained if their peak biofilm biomass reached ≥ 15% of the per-plate WT median biofilm biomass, with the biofilm-deficient mutant Δ*vpsL* exempted.

To assess the predictive signal of individual features, each was evaluated alone using its full feature time-course as input (timepoints 0–30 hours for biofilm biomass, whole-image features, and colony-segmentation derived features; Fig. S2A-D), under the identical random-forest and GroupKFold (5 folds × 5 repeats) protocol, ranked by mean balanced accuracy. To localize predictive signal in time, classifiers were trained on the full feature set at each individual frame (0–30 hours) under the same protocol, and per-frame balanced accuracy was plotted against the full time-course classifier (mean ± s.d. across repeats; Fig. S2E).

### Extension to Other Organisms

The *K. pneumoniae* (Fig. 5B) and multispecies (Fig. 5C) classifiers used the same random-forest configuration and stacked-CLS-embedding representation. For *K. pneumoniae* (five strains), because each plate contained a single strain, plate-grouped CV is not possible; we instead used RepeatedStratifiedKFold over wells (5 folds × 5 repeats), which precludes plate-level batch control. Wells were retained if peak biofilm biomass reached ≥ 15% of the median WT peak biofilm biomass, with the non-biofilm-forming Δ*mrkA* exempted. For the multispecies panel, classification was performed using GroupKFold by plate as above. Balanced accuracy and averaged row-normalized confusion matrices were computed as above. No biofilm biomass filter was applied to exclude timelapses in the multispecies classification analysis.

### Genome-Wide Screen: Mutant Selection and Phenotypic Classification

Biofilm biomass time-series trajectories were normalized to the median peak biofilm biomass of eight WT control wells, where peak biofilm biomass per WT well was defined as the maximum biomass value across all timepoints. The WT reference values were drawn from the defined mutant dataset (Fig. 1 and 2) imaged under identical conditions, providing a stable normalization baseline independent of transposon plate-to-plate variation. Wells in which normalized biofilm biomass exceeded 0.15 at t = 5 hours were excluded as early-onset anomalies inconsistent with normal *V. cholerae* growth and biofilm formation dynamics.

Transposon mutants were assigned to phenotypic classes based on two summary statistics: normalized peak biofilm biomass, the maximal biofilm biomass formed across the growth cycle; and normalized biofilm biomass at t = 25 hours, capturing the persistence of biofilms beyond the point at which WT *V. cholerae* naturally disperses. Classification boundaries were derived from the WT replicate distribution. The upper boundary (B_max) was defined as the maximum WT peak biofilm biomass divided by the mean WT peak biofilm biomass, and the lower boundary (B_min) as the minimum WT peak biofilm biomass divided by the mean WT biofilm biomass, minus 0.25 (providing a conservative margin below the observed WT floor). Mutants were classified as “High Biofilm” if peak biomass exceeded B_max, “Low Biofilm” if peak biomass fell below B_min, “Dispersal Defect” if late-phase biomass exceeded the maximum WT late-phase value while peak biomass remained within [B_min, B_max], and “Normal” otherwise.

### Reimaging Dataset Construction and Quality Filtering

157 mutants classified in the transposon screen as “High Biofilm” or “Dispersal Defect” were selected for reimaging. These strains were re-arrayed into 96-well plates, and *N* = 25 replicates were captured for each strain. After the growth filter, any mutant represented by fewer than five surviving replicate wells was excluded entirely from the analysis.

### Hierarchical Clustering and Heatmap Construction

Hierarchical clustering used the full suite of µPULLI-I features (Table S1). Missing values, which occurred for colony-level features in the absence of microcolonies (e.g. in the Δ*vpsL* strain), were set to zero and zero-variance columns removed. Each replicate well was represented by its full feature-timepoint vector; these vectors were standardized (StandardScaler) and projected onto 50 principal components (PCA, random_state = 0). For each mutant, the median of its replicate wells in PC space defined a phenotypic centroid, and Ward linkage was applied to the pairwise Euclidean distances between centroids (SciPy linkage, method = ‘ward’). This was applied independently to the full reimaging atlas (157 transposon mutants; Fig. S5, Movie S3) and to a functionally annotated subset—mutants from six manually curated functional categories (Motility, O-Antigen Biosynthesis, Polyamine Import, Biotin Biosynthesis, Pyruvate Flux, Vibriobactin Biosynthesis) plus WT (Fig. 3D, Movie S3).

For the static peak-frame heatmap (Fig. 3D), the displayed matrix used, per mutant, the median feature value across replicate wells at that mutant’s peak-biofilm biomass timepoint; values were z-scored across mutants (per feature), clipped to ± 3 standard deviations, and rendered with the RdBu_r diverging colormap. Per-frame versions animated time series over the imaging period (Movie S3).

To visualize each mutant’s phenotypic profile, we constructed feature-by-mutant barcode heatmaps in which each column is a mutant (or condition) summarized by the median of its replicate wells at that mutant’s peak-biomass frame, and each row is a feature. Two color scales were used depending on the comparison. For the *V. cholerae* training set (Fig. 2C) and the reimaging atlas (Fig. 3D), values were z-scored across mutants per feature. For the clean-deletion (Fig. 4C) and compound-treatment (Fig. 5A) comparisons, values were expressed relative to WT as 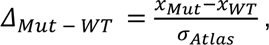 where *x* is the feature value at peak biofilm biomass, and *σ*_*Atlas*_ is the standard deviation of that same feature across the transposon reimaging atlas (each atlas mutant feature value taken at its own peak-biomass frame; Fig. S5). In all cases values were clipped to ±3 and rendered with the RdBu_r diverging colormap.

### Statistical Analysis of Transposon Reimaging Replicate Consistency

To test whether each mutant’s measured phenotype is reproducible across replicate wells, distances between imaging replicates in quantitative feature space (including biofilm biomass, colony segmentation-derived features, and whole-image features at timepoints 9-30 h) were compared for replicates of the same mutant versus replicates of different mutants (Fig. S5B). Features were standardized across replicates and used directly, without dimensionality reduction. The dataset comprises 3,867 imaging replicates across 157 mutants and WT (median 24 per mutant, range 6–24). Euclidean distances were computed between all 6,728,946 possible pairs of replicates, and pairs were partitioned into same-mutant (41,110) and different-mutant (6,687,836) sets. The null hypothesis throughout is that mutant identity carries no information about distance, such that two replicates of the same mutant are no more similar than two replicates of different mutants. It was tested by permuting mutant labels (10,000 permutations); permutation p values are bounded below by 1/(permutations + 1) and are reported as p < 0.0001. Two null models were used: labels shuffled across all 157 mutants, and labels shuffled only among mutants imaged on the same plate, the latter holding plate composition fixed. Across all pairs of replicates, the same-mutant and different-mutant distance distributions were compared directly to ask how reliably a replicate pair can be told from an unrelated one. Their separation was summarized as the area under the ROC curve—equivalently the normalized Mann–Whitney U—the probability that a randomly chosen same-mutant pair is closer than a randomly chosen different-mutant pair (Fig. S5B, left). At the level of individual mutants, to establish that reproducibility is general rather than driven by a subset, each mutant’s mean distance to its own replicates was compared with the distribution that same quantity takes under permutation, pooled across all 157 mutants (Fig. S5B, right). Finally, as a check that batch structure does not account for the separation, different-mutant distances were compared for replicate pairs imaged on the same plate versus on different plates.

### Generative AI

The authors made use of large language models (Claude and Gemini) to support manuscript editing and code development. All outputs were critically reviewed and revised by the authors, who take full responsibility for the content of this work.

## DATA AVAILABILITY STATEMENT

The tabular data underlying all figures, along with liquid handling protocols, are available on KiltHub (DOI: 10.1184/R1/33333711). RNA sequencing data in this study have been deposited in the NCBI Sequence Read Archive (SRA) under BioProject accession number PRJNA1513060. Image datasets, including raw, processed, and masked µPULLI images are available on the EMBL-EBI BioImage Archive (DOI: 10.6019/S-BIAD3830).

## CODE AVAILABILITY STATEMENT

Scripts for image analysis, including graphical user interfaces for µPULLI-DL and µPULLI-I, were written in Python (v3.9) and are publicly available on GitHub (https://github.com/BridgesLabCMU/uPULLI-I and https://github.com/BridgesLabCMU/uPULLI-DL), along with readme instructions. Scripts for data processing, tabular information, and figure generation were written in Python (v3.11) and are available at (https://github.com/BridgesLabCMU/uPULLI-figures/).

## DECLARATION OF INTERESTS

The authors declare no competing interests.

## AUTHOR CONTRIBUTIONS

A.A.B. conceived of the project. A.A.B., S.S.M., J.J.D., D.B., G.C., M.B., S.G., S.T., and I.V.M. designed experiments and analyses. S.S.M. developed the µPULLI-DL and µPULLI-I analysis pipelines and wrote the analysis software. A.A.B., S.S.M., J.J.D., D.B., G.C., M.B., S.G., S.T., and I.V.M. performed experiments and analyzed data. I.V.M., J.J.D., S.G., D.J.S., O.C. and L.A.M. provided strains, reagents, and computational resources. A.A.B., S.S.M., and J.J.D. prepared the figures. A.A.B., S.S.M., and J.J.D. were responsible for data curation and wrote the manuscript. All authors reviewed and edited the manuscript. A.A.B., L.A.M., and O.C. supervised the work and acquired funding. A.A.B. coordinated and administered the project.

## ACKNOWLEDGEMENTS

We would like to thank members of the Bridges lab for feedback throughout the manuscript research and writing process. We thank Dr. Christopher Waters for sharing the *V. cholerae* transposon library, Dr. Vaughn Cooper for sharing *A. baumannii* strains, Dr. Catherine Armbruster for *P. aeruginosa*, Dr. Courtney Ellison for *A. baylyi*, Dr. Tony Richardson for *S. aureus*, and Dr. Daria van Tyne for *E. faecalis*. This work was supported by NIAID grant R00AI158939, NIGMS grant 1R35GM160020, a Shurl and Kay Curci Foundation grant (https://curcifoundation.org/), a Kaufman Foundation New Investigator Research Grant KA2023-136488 (https://kaufman.pittsburghfoundation.org/), a Damon Runyon Cancer Research Foundation Dale F. Frey Award for Breakthrough Scientists 2302-17 (https://www.damonrunyon.org/), and startup funds from Carnegie Mellon University to A.A.B., seed funding through the Carnegie Mellon University AI4Bio program to A.A.B. and O.C., and an NIGMS grant R35GM150588 and a Samuel and Emma Winters Foundation grant to L.A.M.

## SUPPLEMENTARY MATERIAL

**Table S1.** Mathematical features extracted using µPULLI-I.

| Feature | Description | Units |
| --- | --- | --- |
| Biofilm Biomass | Total segmented biofilm signal in the field of view | a.u. |
| **Colony-segmentation-derived features** |  |  |
| Colonies | Number of segmented colonies in the field of view | count |
| Area | Mean colony cross-sectional area | $\mu\text{m}^2$ |
| Area Variability | Standard deviation of colony area across colonies in the field of view | $\mu\text{m}^2$ |
| Eccentricity | Mean colony eccentricity (0 = circle, → 1 = elongated) | dimensionless |
| Major Axis Length | Mean length of the longest colony axis | $\mu\text{m}$ |
| Intensity | Mean brightfield pixel intensity inside colonies | a.u. |
| Intensity Kurtosis | Kurtosis of the per-colony mean-intensity distribution | dimensionless |
| Intensity Variability | Coefficient of variation of background pixel intensity | dimensionless |
| Radial Offset | Mean distance from each colony centroid to the center of the field of view | $\mu\text{m}$ |
| Max Distance | Longest edge in the minimum spanning tree over colony centroids | $\mu\text{m}$ |
| Nearest Neighbor | Mean nearest-neighbor distance between colony centroids | $\mu\text{m}$ |
| NN Variability | Standard deviation of nearest-neighbor distance | $\mu\text{m}$ |
| **Whole-image texture features (Haralick GLCM + entropy)** |  |  |
| Global Entropy | Shannon entropy of the intensity histogram in the field of view | bits |
| Energy | Angular second moment of the gray-level co-occurrence matrix | dimensionless |
| Contrast | GLCM contrast (local intensity variation) | intensity<sup>2</sup> |
| Correlation | Linear dependency of gray levels in the GLCM | dimensionless |
| Variance | GLCM sum of squares (intensity variance) | intensity<sup>2</sup> |
| Inv Diff Moment | Inverse difference moment (image homogeneity) | dimensionless |
| Sum Average | Mean of the GLCM sum distribution | intensity |
| Sum Variance | Variance of the GLCM sum distribution | intensity<sup>2</sup> |
| Sum Entropy | Entropy of the GLCM sum distribution | bits |
| Entropy | Entropy of the full GLCM | bits |
| Diff Variance | Variance of the GLCM difference distribution | intensity<sup>2</sup> |
| Diff Entropy | Entropy of the GLCM difference distribution | bits |
| IMC 1 | Information measure of correlation 1 | dimensionless |
| IMC 2 | Information measure of correlation 2 | dimensionless |

Units of “a.u.” (arbitrary units) refer to camera-intensity values on the unprocessed brightfield image. a.u. = arbitrary units; GLCM = gray level co-occurrence matrix.

**Table S2.**
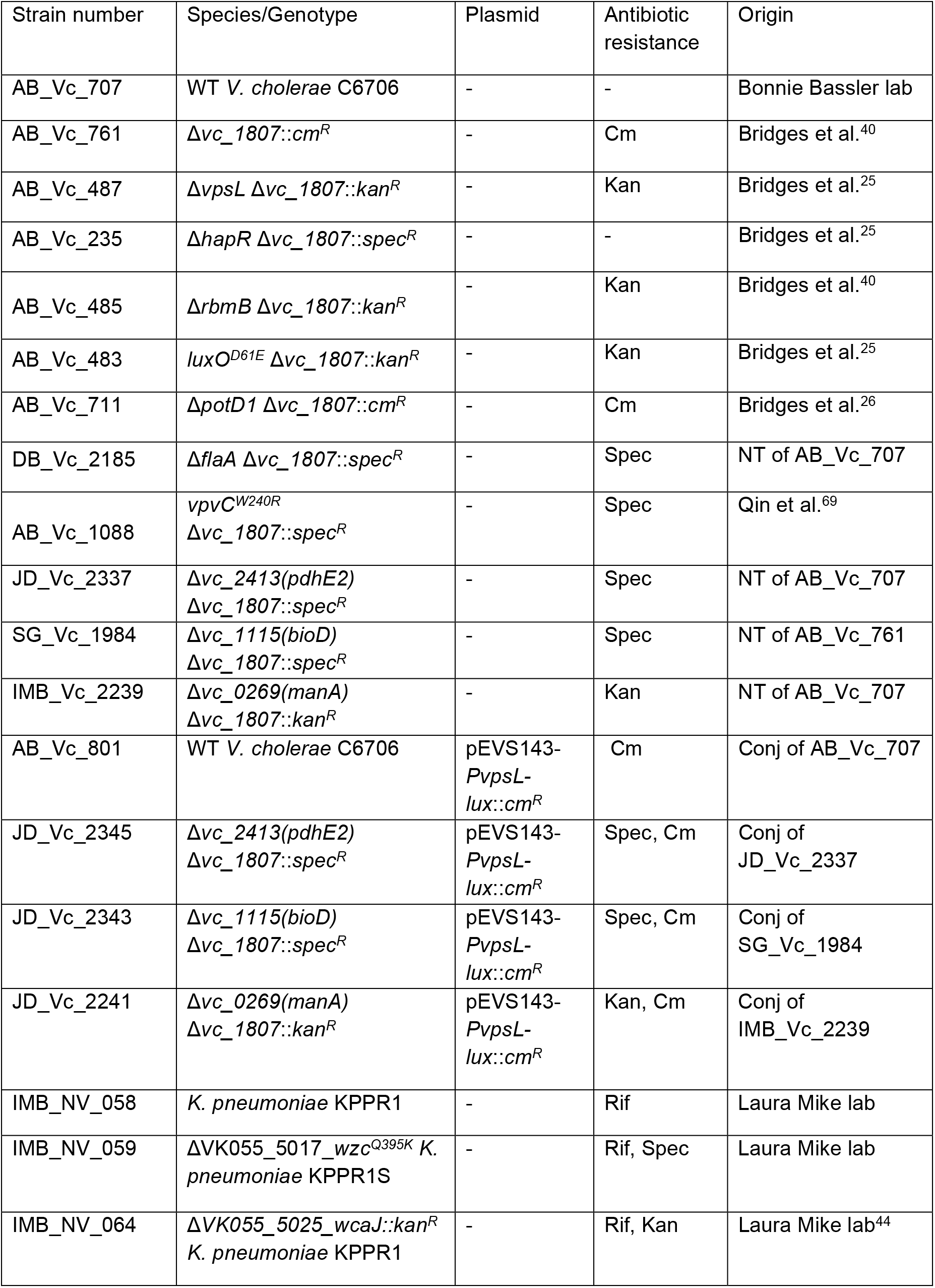
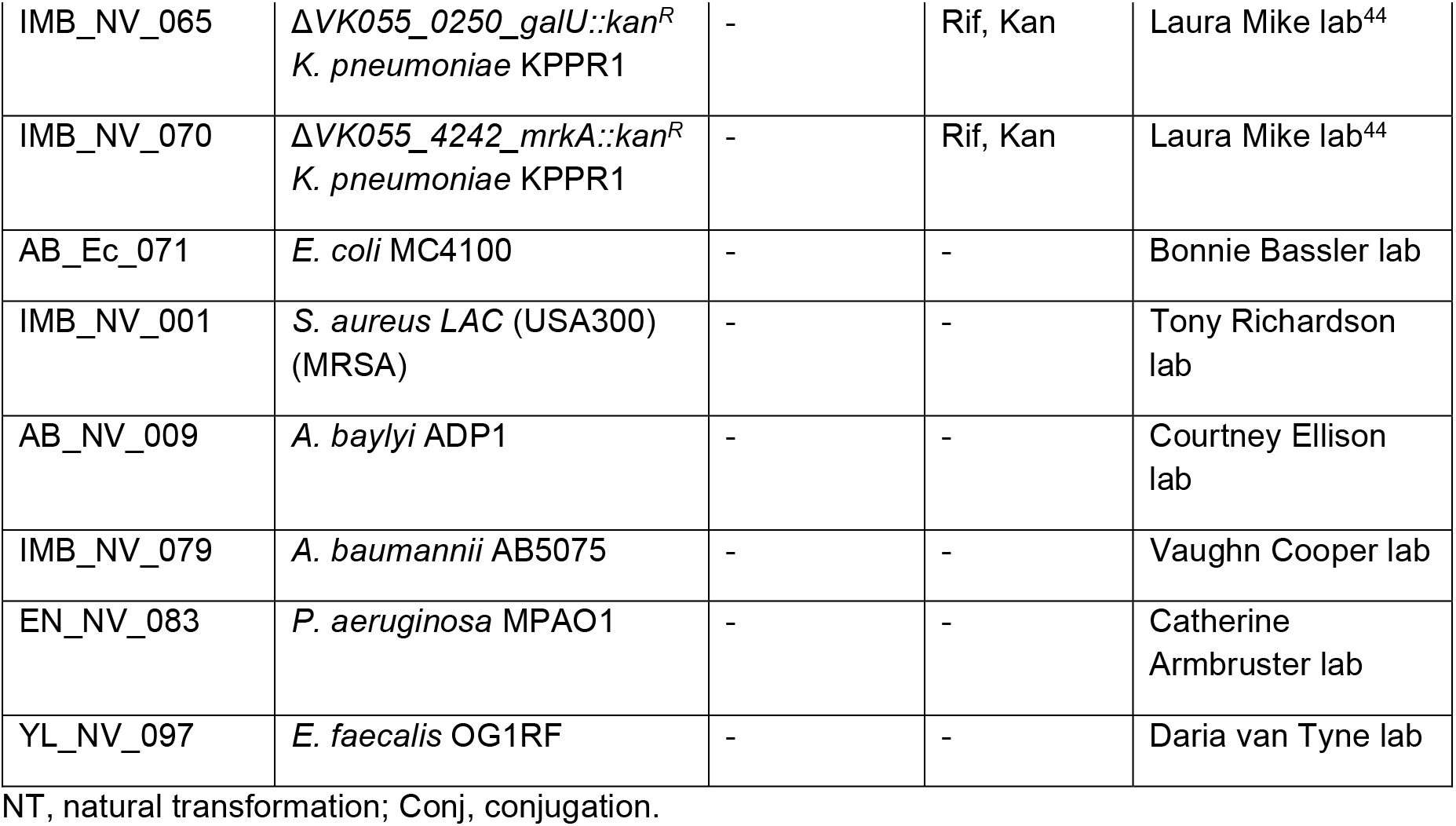
Strains used in this study.

| Strain number | Species/Genotype | Plasmid | Antibiotic resistance | Origin |
| --- | --- | --- | --- | --- |
| AB_Vc_707 | WT *V. cholerae* C6706 | - | - | Bonnie Bassler lab |
| AB_Vc_761 | $\Delta vc\_1807::cm^R$ | - | Cm | Bridges et al.<sup>40</sup> |
| AB_Vc_487 | $\Delta vpsL \Delta vc\_1807::kan^R$ | - | Kan | Bridges et al.<sup>25</sup> |
| AB_Vc_235 | $\Delta hapR \Delta vc\_1807::spec^R$ | - | - | Bridges et al.<sup>25</sup> |
| AB_Vc_485 | $\Delta rbmB \Delta vc\_1807::kan^R$ | - | Kan | Bridges et al.<sup>40</sup> |
| AB_Vc_483 | $luxO^{D61E} \Delta vc\_1807::kan^R$ | - | Kan | Bridges et al.<sup>25</sup> |
| AB_Vc_711 | $\Delta potD1 \Delta vc\_1807::cm^R$ | - | Cm | Bridges et al.<sup>26</sup> |
| DB_Vc_2185 | $\Delta flaA \Delta vc\_1807::spec^R$ | - | Spec | NT of AB_Vc_707 |
| AB_Vc_1088 | $vpvC^{W240R} \Delta vc\_1807::spec^R$ | - | Spec | Qin et al.<sup>69</sup> |
| JD_Vc_2337 | $\Delta vc\_2413(pdxE2) \Delta vc\_1807::spec^R$ | - | Spec | NT of AB_Vc_707 |
| SG_Vc_1984 | $\Delta vc\_1115(bioD) \Delta vc\_1807::spec^R$ | - | Spec | NT of AB_Vc_761 |
| IMB_Vc_2239 | $\Delta vc\_0269(manA) \Delta vc\_1807::kan^R$ | - | Kan | NT of AB_Vc_707 |
| AB_Vc_801 | WT *V. cholerae* C6706 | pEVS143-*PvpsL-lux::cm^R* | Cm | Conj of AB_Vc_707 |
| JD_Vc_2345 | $\Delta vc\_2413(pdxE2) \Delta vc\_1807::spec^R$ | pEVS143-*PvpsL-lux::cm^R* | Spec, Cm | Conj of JD_Vc_2337 |
| JD_Vc_2343 | $\Delta vc\_1115(bioD) \Delta vc\_1807::spec^R$ | pEVS143-*PvpsL-lux::cm^R* | Spec, Cm | Conj of SG_Vc_1984 |
| JD_Vc_2241 | $\Delta vc\_0269(manA) \Delta vc\_1807::kan^R$ | pEVS143-*PvpsL-lux::cm^R* | Kan, Cm | Conj of IMB_Vc_2239 |
| IMB_NV_058 | *K. pneumoniae* KPPR1 | - | Rif | Laura Mike lab |
| IMB_NV_059 | $\Delta VK055\_5017\_wzc^{Q395K}$ *K. pneumoniae* KPPR1S | - | Rif, Spec | Laura Mike lab |
| IMB_NV_064 | $\Delta VK055\_5025\_wcaJ::kan^R$ *K. pneumoniae* KPPR1 | - | Rif, Kan | Laura Mike lab<sup>44</sup> |
| IMB_NV_065 | ΔVK055_0250_galU::kan<sup>R</sup><br>*K. pneumoniae* KPPR1 | - | Rif, Kan | Laura Mike lab<sup>44</sup> |
| IMB_NV_070 | ΔVK055_4242_mrkA::kan<sup>R</sup><br>*K. pneumoniae* KPPR1 | - | Rif, Kan | Laura Mike lab<sup>44</sup> |
| AB_Ec_071 | *E. coli* MC4100 | - | - | Bonnie Bassler lab |
| IMB_NV_001 | *S. aureus* LAC (USA300)<br>(MRSA) | - | - | Tony Richardson<br>lab |
| AB_NV_009 | *A. baylyi* ADP1 | - | - | Courtney Ellison<br>lab |
| IMB_NV_079 | *A. baumannii* AB5075 | - | - | Vaughn Cooper lab |
| EN_NV_083 | *P. aeruginosa* MPAO1 | - | - | Catherine<br>Armbruster lab |
| YL_NV_097 | *E. faecalis* OG1RF | - | - | Daria van Tyne lab |

NT, natural transformation; Conj, conjugation.

**Table S3.**
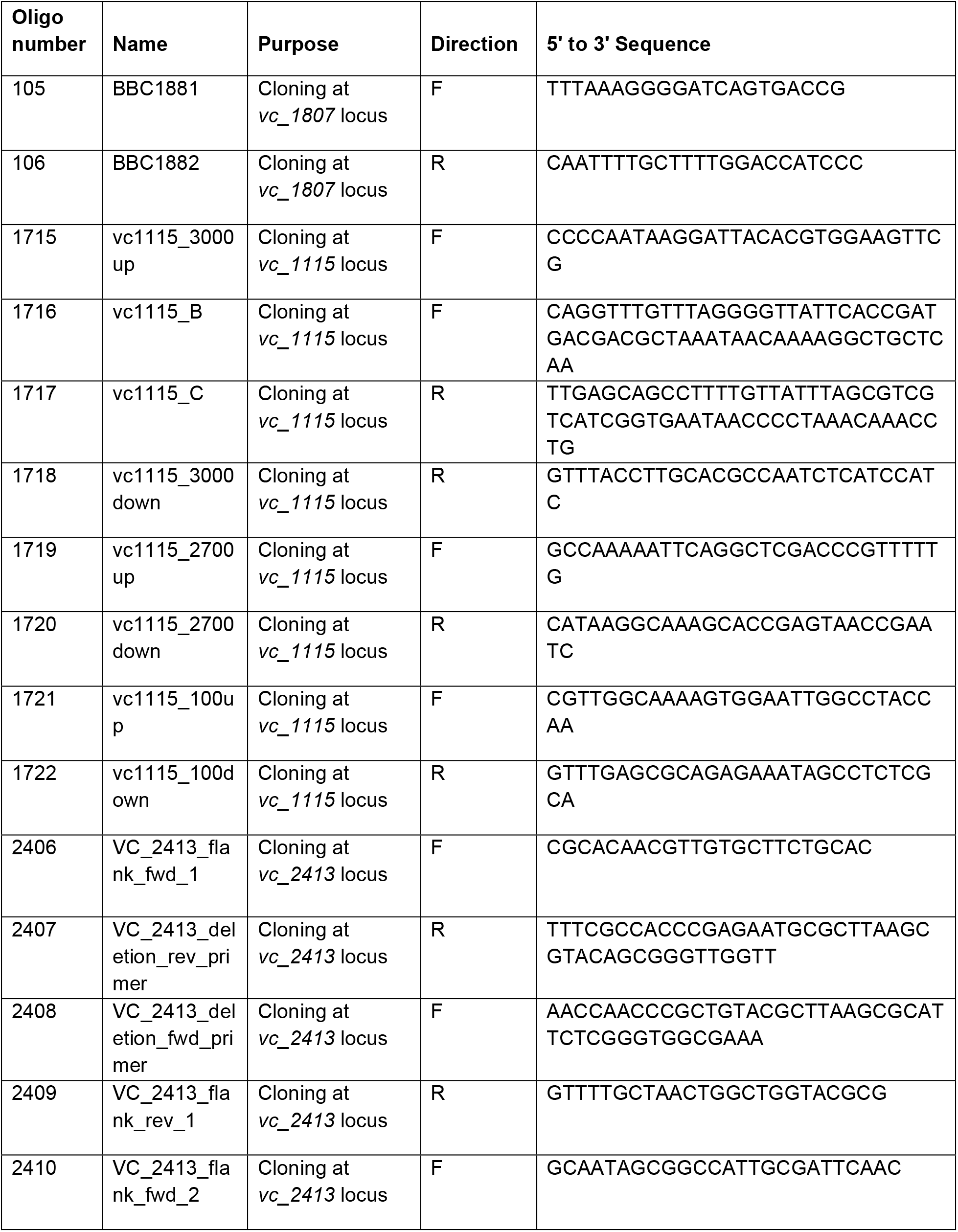
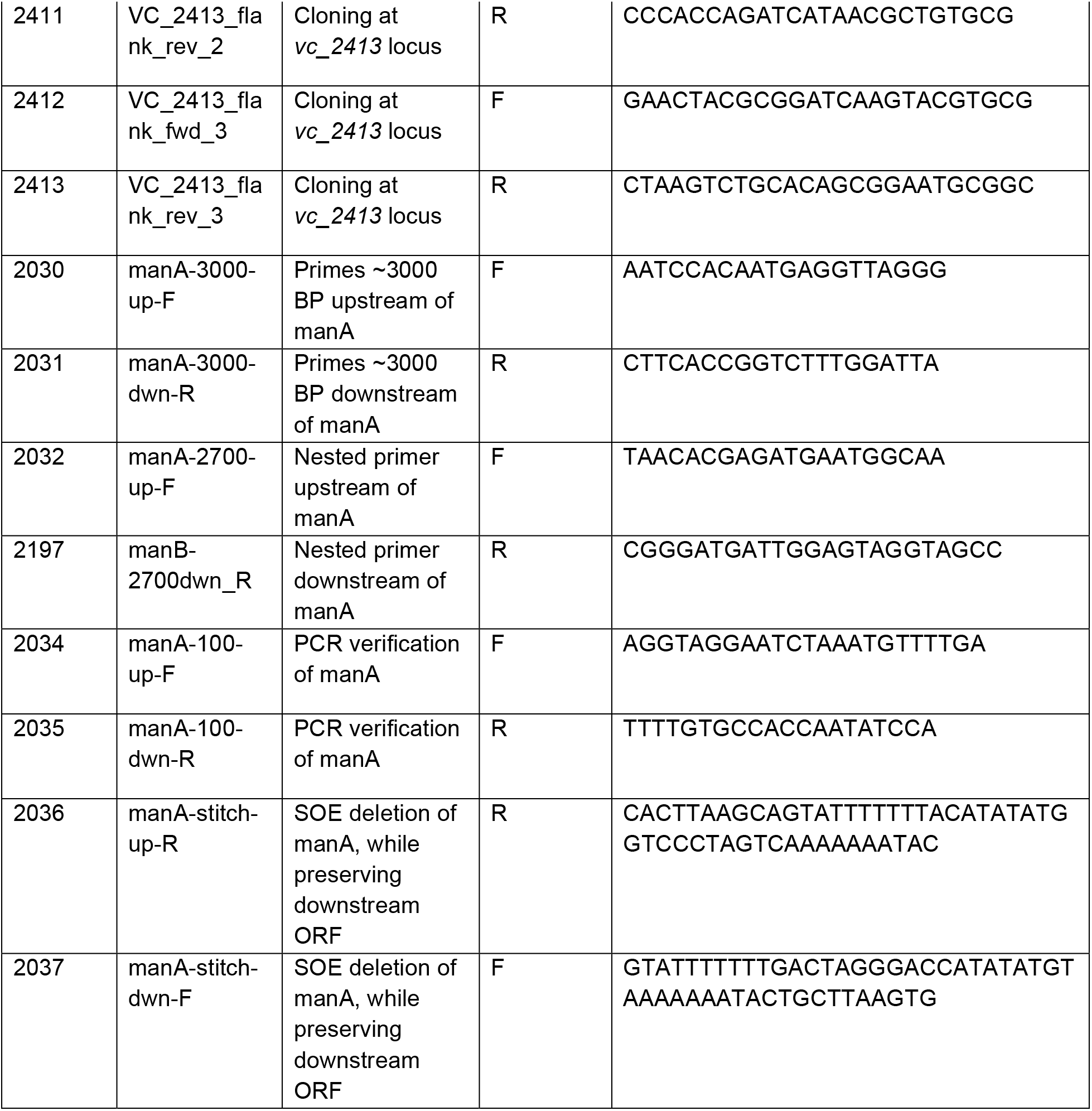
Oligos used in this study.

**Figure S1.**
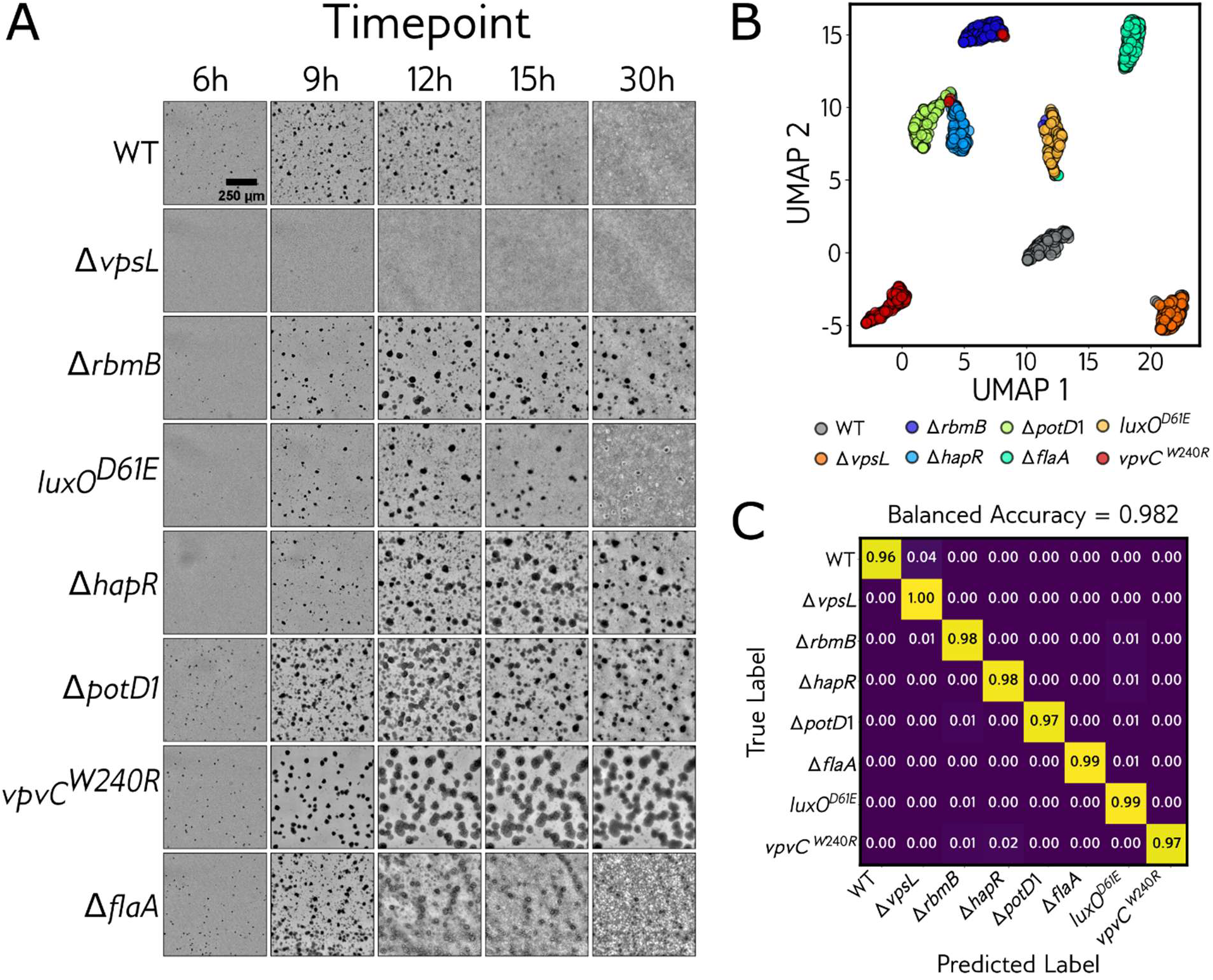
DINOv2 patch-token embeddings of label-free brightfield timelapse imaging distinguish eight *V. cholerae* mutants. **(A)** Representative frames from 0–30 hours of 10× brightfield imaging of culture growth reveals qualitatively distinct colony formation, aggregation, dispersal, and morphological dynamics across the eight indicated strains. **(B)** UMAP dimensionality reduction of DINOv2 patch-token features (nine per-frame regional patch descriptors averaged to one 768-dimensional descriptor and concatenated across the timecourse; see Methods), where each point denotes an individual timelapse replicate and point colors denote distinct *V. cholerae* strain identity. **(C)** Confusion matrix for a random-forest classifier trained on the patch-token embeddings, with performance assessed by plate-grouped cross-validation (5 folds × 5 repeats, wells from the same plate never split across train and test). Cells in the matrix are colored by classification accuracy; the model achieved 98.2% balanced classification accuracy across all eight strain labels, comparable to the CLS-token representation (Fig. 1E). *N* = 1,149 growth-filtered replicate wells.

**Figure S2.**
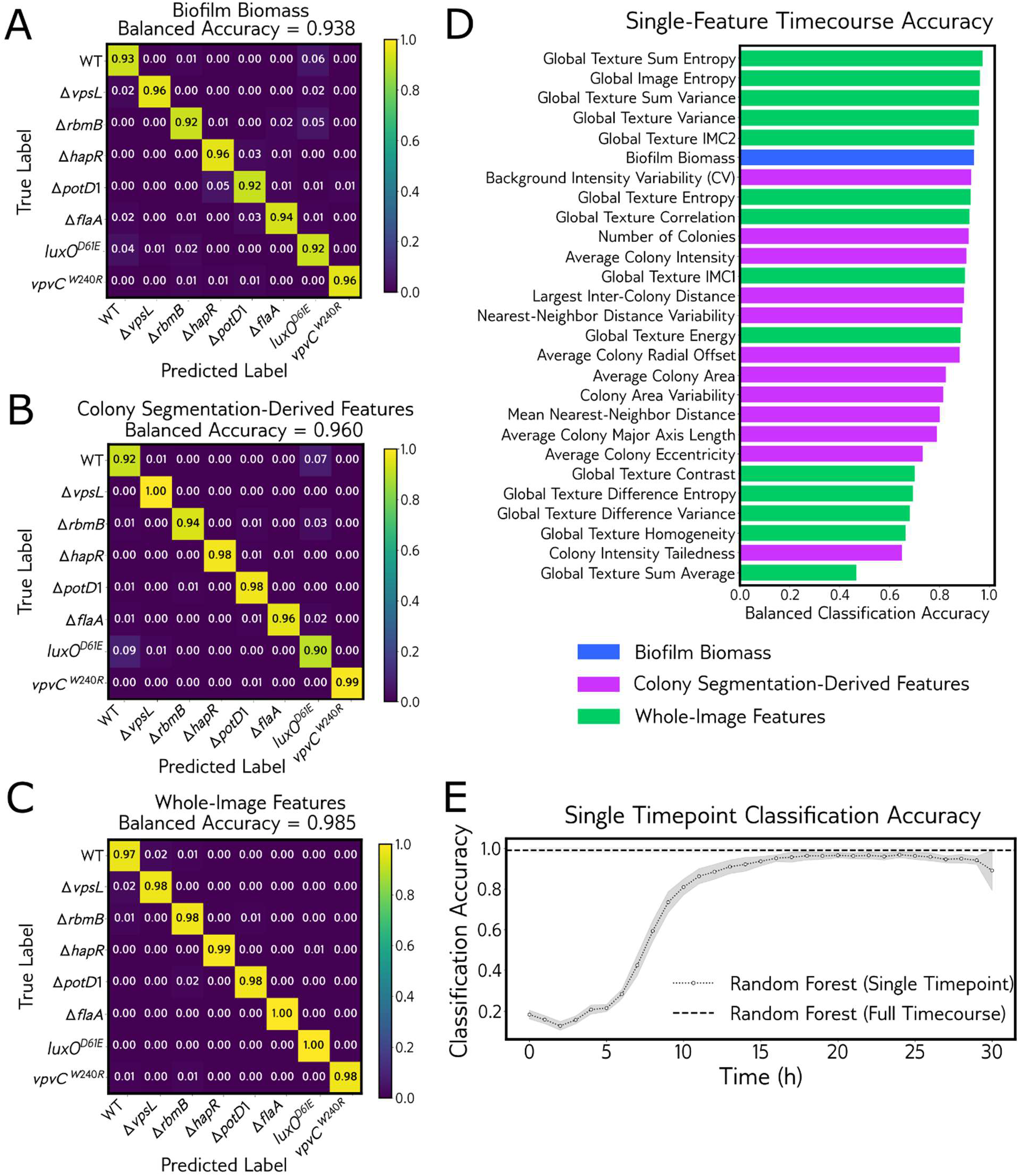
Feature-class and temporal decomposition of interpretable genotype classification. Companion to Fig. 2, dissecting which quantitative feature classes and which timepoints drive random-forest classification of the eight *V. cholerae* strains. All panels use the cross-validation scheme of Fig. 2 (random forest, GroupKFold with plates as groups, 5 folds × 5 repeats, balanced accuracy). (A–C) Confusion matrices for classifiers trained on a single feature class in isolation: **(A)** biofilm biomass trajectories alone (93.8% balanced accuracy), **(B)** segmented-microcolony features alone (96.0%), and **(C)** whole-image texture features alone (98.5%). Color bars indicate per-class classification accuracy. **(D)** Balanced classification accuracy of each individual feature when trained on that feature’s full timecourse alone, ranked from most to least discriminative; bars are colored by feature class (biofilm biomass, whole-image texture, colony-segmentation-derived). **(E)** Mutant separability across time. Dotted line: balanced accuracy of a random forest trained on all features at a single timepoint, plotted against imaging time (mean ± s.d. across cross-validation folds); dashed line: the full-timecourse baseline trained on all timepoints together (99.0% ± 1.3%). Separability is near chance in early culture growth, rises steeply between ∼5 and ∼12 hours as spatial structure emerges, and plateaus near the full-timecourse accuracy for the remainder of the timecourse. *N* = 1,149 growth-filtered replicate wells.

**Figure S3.**
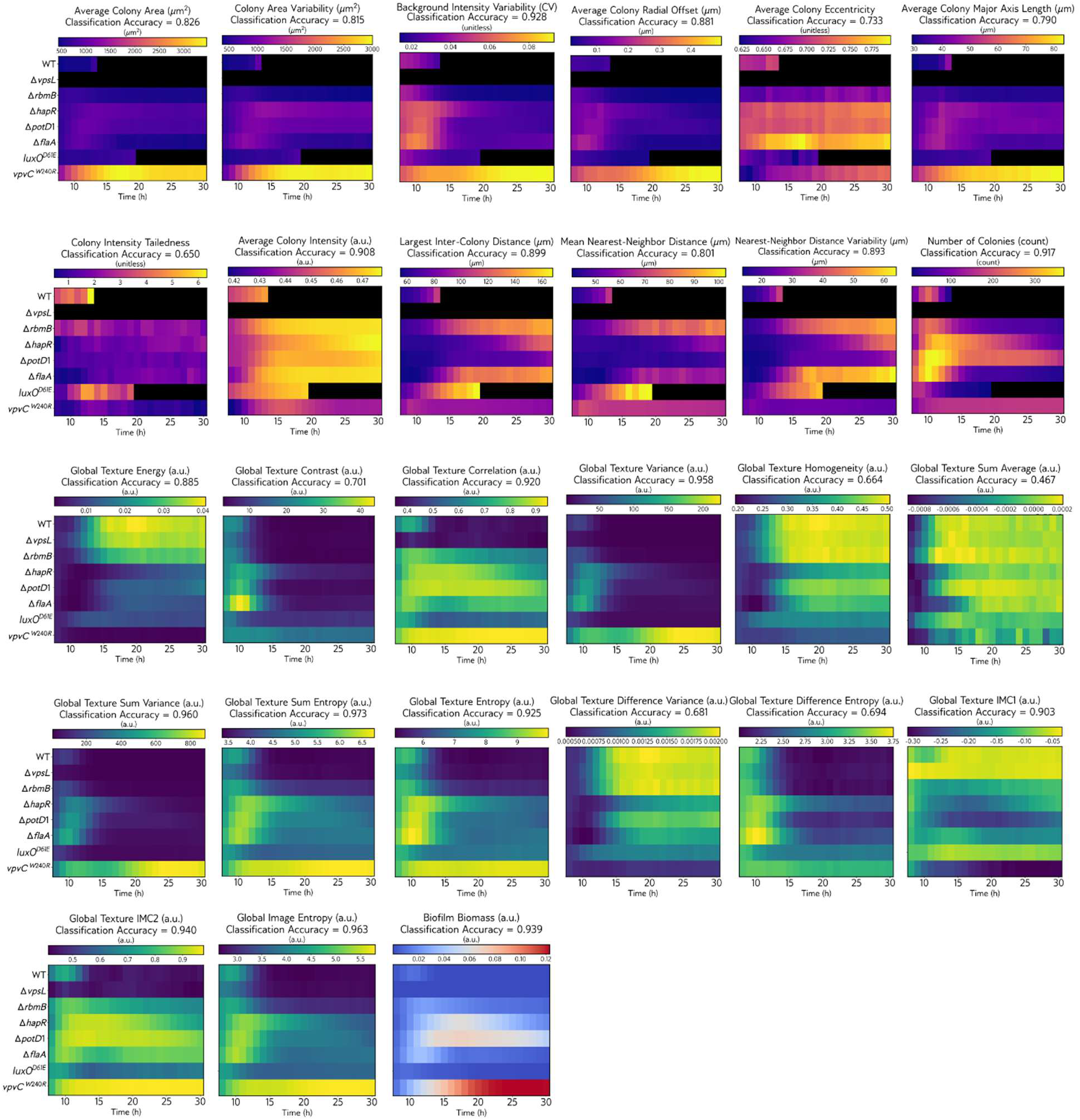
The complete set of feature-class temporal heatmaps summarized in Fig. 2F. The single-feature random-forest balanced classification accuracy (Fig. S2D) is shown above each panel. Plasma colormap denotes colony segmentation-derived features, viridis colormap denotes whole-image and Haralick features, and blue-to-red (RdBu_r) colormap denotes biofilm biomass trajectories. Black indicates absence of colony segmentation-derived features due to lack of colonies detected in the images.

**Figure S4.**
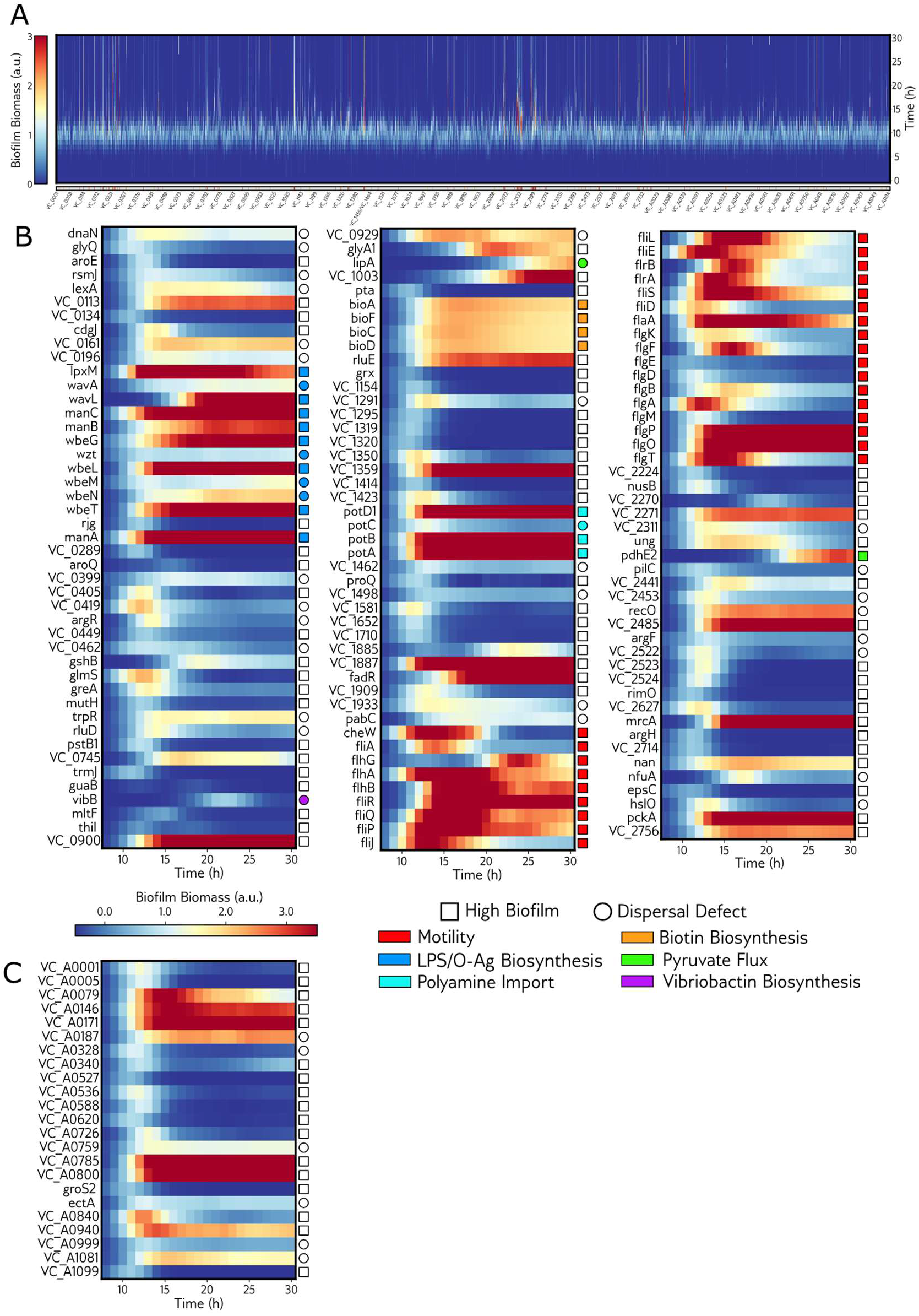
Genome-wide transposon biomass screen of *V. cholerae* biofilm formation. Biofilm-biomass trajectories for an ordered transposon library (one insertion per non-essential gene; 2,850 mutants), normalized to the WT peak mean and classified against the WT replicate distribution as High Biofilm, Low Biofilm, Dispersal Defect, or Normal (see Methods). **(A)** Genome-wide heatmap: each column is a mutant, ordered by chromosomal locus (Chromosome I, VC_####, then Chromosome II, VC_A####); rows are imaging time; color is normalized biofilm biomass. The strip along the bottom marks the mutants selected for reimaging (High Biofilm or Dispersal Defect). (B, C) Biofilm biomass trajectories for the 157 transposon mutants selected for reimaging, shown per gene, split by chromosome—**(B)** Chromosome I (134 mutants) and **(C)** Chromosome II (23 mutants)—one row per mutant, one column per hour (8–30 hours). Rows are labeled by gene name (or locus number where unnamed); the marker at right gives phenotype (square, High Biofilm; circle, Dispersal Defect) and functional annotation (fill color; open marker, none).

**Figure S5.**
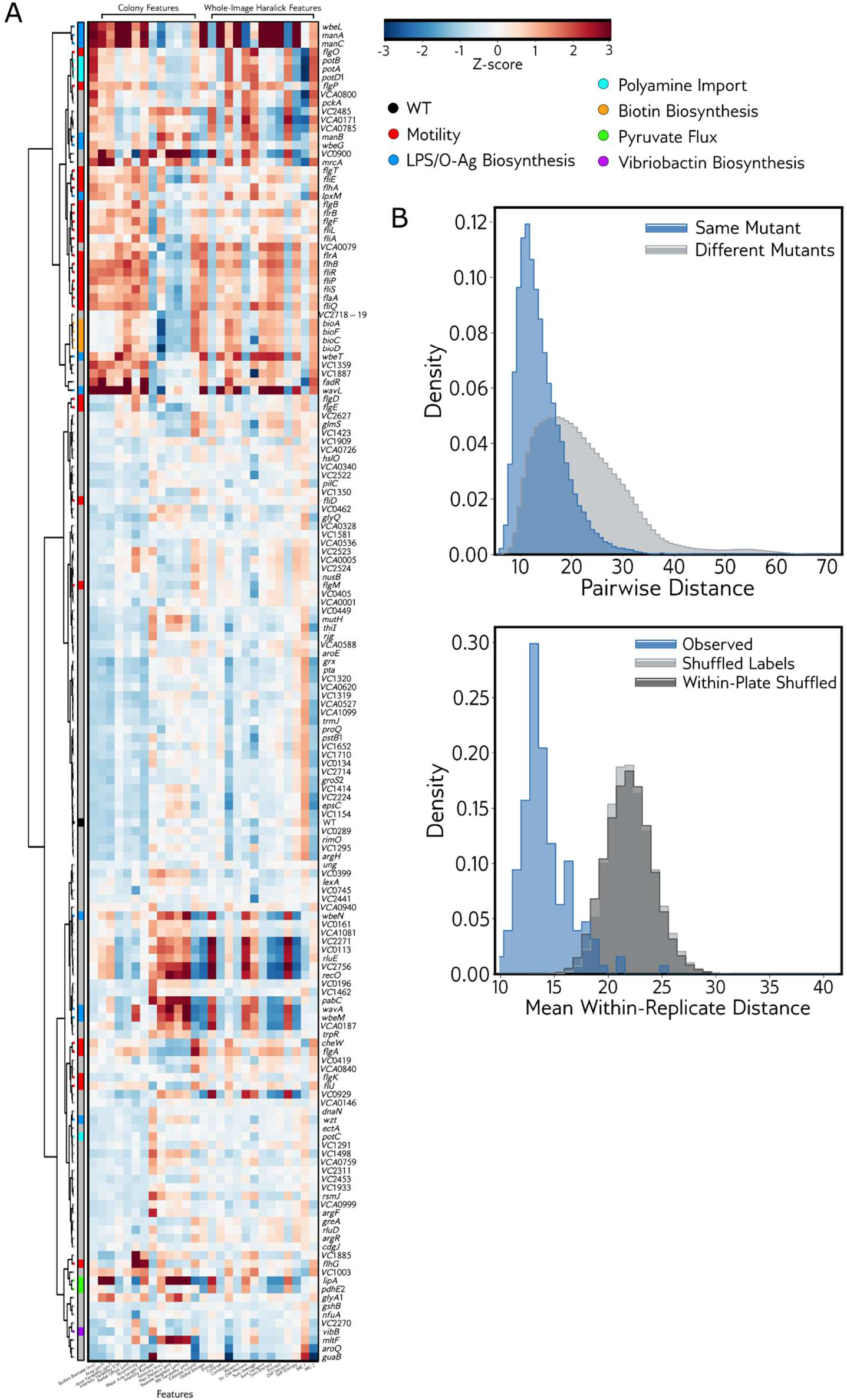
Structure and reproducibility of the transposon reimaging atlas from quantitative features. **(A)** All 157 reimaging mutants clustered by their quantitative features, each mutant summarized as the centroid of its replicate wells in PCA-50 space (Ward linkage; Methods). PCA-50 space was constructed from an original dataset including 22 timepoints (9-30 h) of biofilm biomass, 14 whole-image, and 12 colony segmentation-derived features. Heatmap shows feature values at each mutant’s peak-biofilm biomass frame, z-scored across all 157 mutants. Color strip, functional annotation colored according to legend and as in Fig. 3 (WT, black; unannotated, gray). **(B)** Reproducibility of each mutant phenotype in the transposon reimaging mutant atlas. The 157 mutants are represented by 3,867 imaging replicates (median 24 per mutant), where each replicate is treated as a data point in phenotypic feature space spanning biofilm biomass, whole-image, and colony-level features (timepoints 9-30 h; Fig. 3B). (B, left) Distribution of pairwise Euclidean distances between same-mutant (blue) and different-mutant (gray) pairs of replicates. Same-mutant pairs are closer (means 14.0 versus 22.1 Euclidean distance; Area under curve (AUC) = 0.81, p < 0.0001; see Methods). (B, right) Distribution across the 157 mutants of each mutant’s mean distance to its own replicates (blue), against the same quantity when mutant labels are shuffled (gray, across the whole atlas; dark gray, only within an imaging plate, controlling for batch). 156 of 157 mutants fall below the null mean (AUC = 0.99, p < 0.0001).

**Figure S6.**
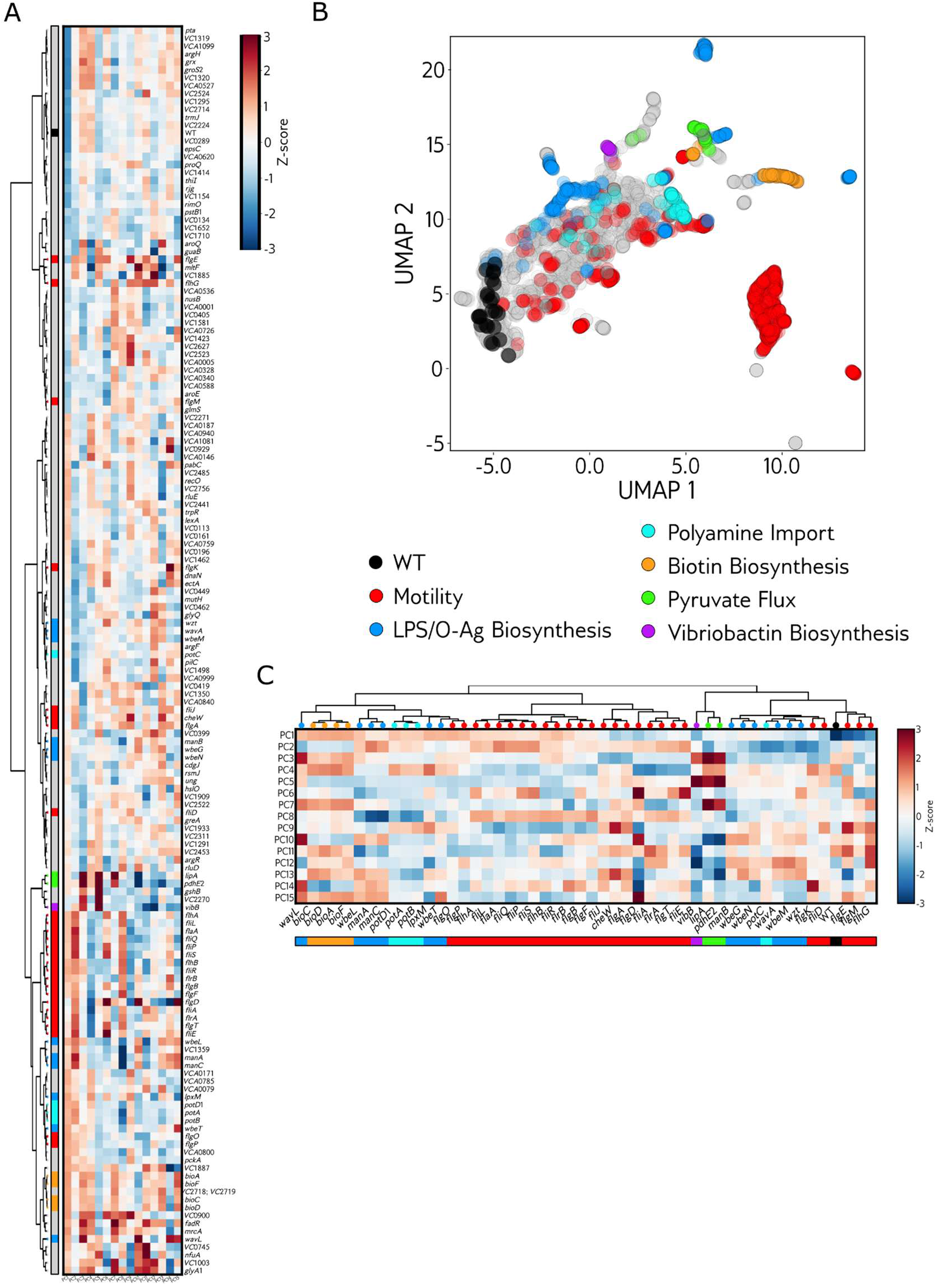
DINOv2-embedding views of the transposon reimaging atlas. The deep-embedding counterparts of the quantitative-feature analyses in Fig. 3 and Fig. S6. **(A)** Dendrogram and heatmap of all 157 reimaging mutants from their DINOv2 embeddings: per-mutant centroids in the embedding space were clustered by Ward linkage on the top 15 principal components (PCs), and the heatmap shows those 15 PCs z-scored across mutants (mutants in dendrogram-leaf order; leaves colored by functional annotation). **(B)** UMAP of the reimaging mutants computed from the DINOv2 embeddings (CLS token, PCA-50; n_neighbors = 10, min_dist = 0.1); each point is a replicate well, colored by functional annotation (WT, black; unclassified, gray). The functional pathways are recovered as distinct regions of the embedding manifold. **(C)** As in (A), but restricted to the functionally-annotated subset (the six highlight pathways + WT; 50 mutants), with a functional-annotation legend. Color scale, Z-score; leaf and strip colors denote functional annotation.

**Figure S7.**
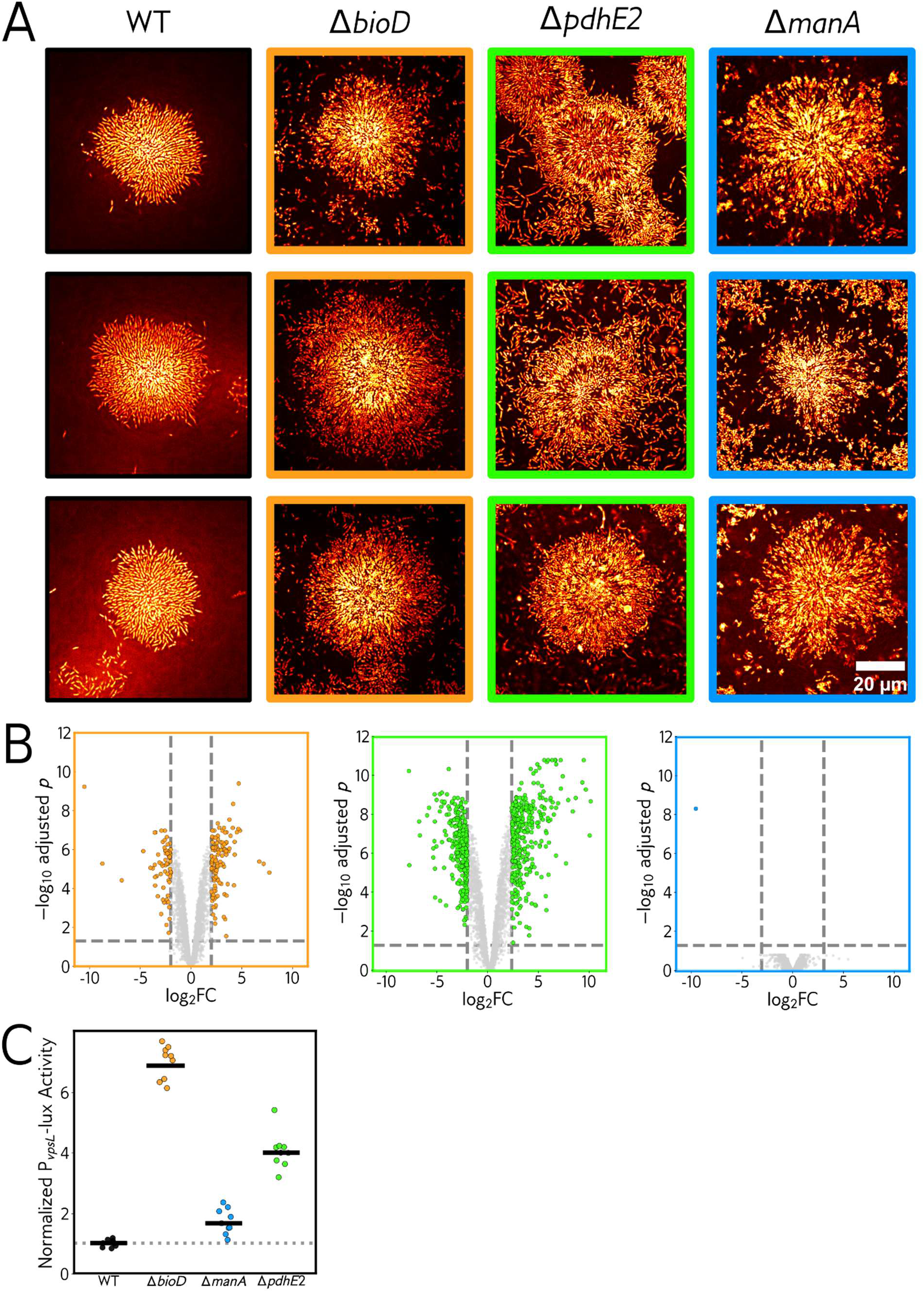
Confocal micrographs and matrix reporter output for mutant strains. **(A)** As in Fig. 4D, replicate cropped high-resolution confocal fluorescence images of representative microcolonies (medial plane) for WT, Δ*bioD*, Δ*pdhE2*, and Δ*manA*. Cells were stained with 500 µg/mL DAPI. Scale bar, 20 µm. **(B)** Volcano plots of RNA-sequencing results for Δ*bioD* (left, orange), Δ*pdhE2* (center, green), and Δ*manA* (right, blue) relative to WT, respectively. |log_2_FC| > 2 and P-value (Benjamini-Hochberg) < 0.05 were used as significance cutoffs (dashed lines, blue points are significant). *N* = 3 biological replicates. **(C)** Quantification of *P_vpsL_-lux* reporter activity for the indicated mutants. Points represent peak luminescence (highest luminescence achieved across the growth cycle for that culture) for individual replicates, normalized to the mean of the WT strain peak (*N* = 3 biological replicates and 3 technical replicates). Crossbars represent mean values.

## SUPPLEMENTARY TEXT

### Interpretation of whole-image Haralick textural statistics

To interpret this texture-based signal, we examined how whole-image features related to the physically defined colony-level features via hierarchical clustering (Fig. 2C). Haralick sum entropy clustered closely with colony segmentation-derived features such as colony area, major axis length, area variability, and radial offset, suggesting that this otherwise abstract texture statistic largely reflects the size and dispersion of microcolonies. Other texture statistics resolve similarly: contrast and difference entropy group with colony eccentricity, IMC1 with colony number and intensity, and correlation and IMC2 with biofilm biomass. Thus, the colony segmentation-derived features provide physical grounding for Haralick features that are not interpretable in isolation, while the whole-image Haralick features offer a segmentation-free complement that remains informative even when strains fail to form microcolonies altogether, a regime where colony-level features cannot be computed at all.

## SUPPLEMENTARY DATA

**Data S1.** Results from initial transposon screen.

## SUPPLEMENTARY MOVIES

**Movie S1.** Brightfield timelapses of the eight *V. cholerae* strains shown in Fig. 1B.

**Movie S2.** Brightfield timelapses of the transposon mutants shown in Fig. 3C, grouped by functional category.

**Movie S3.** Animation of the reimaging-atlas dendrogram, showing how each feature class evolves across the developmental timecourse.

**Movie S4.** Brightfield timelapses of the small-molecule conditions shown in Fig. 5A.

**Movie S5.** Brightfield timelapses of the five *K. pneumoniae* strains shown in Fig. 5B.

**Movie S6.** Brightfield timelapses of the eight bacterial species shown in Fig. 5C.

## INTERACTIVE PLOTS

**Interactive Plot 1.** Interactive UMAP of the transposon reimaging landscape based on quantitative features extracted using µPULLI-I. Clicking any replicate displays its peak biofilm biomass image.

**Interactive Plot 2.** Interactive UMAP of the transposon reimaging landscape based on PCA-50 dimensionality reduction of CLS token embeddings extracted from the encoder layer of DINOv2 using µPULLI-DL. Clicking any replicate displays its peak biofilm biomass image.

**Interactive Plot 3.** Interactive dendrogram and heatmaps of the transposon reimaging atlas at all timepoints (0-30 h). Clicking any mutant displays representative images from three replicates.

**Interactive Plot 4.** Interactive volcano plot of RNA-sequencing results for Δ*bioD* relative to WT. A Log_2_fold > 2 and P-value (Benjamini-Hochberg) < 0.05 were used as significance cutoffs (dashed lines, blue points are significant). *N* = 3 biological replicates. Selecting a functional category on the left-hand panel highlights associated genes in the volcano plot.

**Interactive Plot 5.** Interactive volcano plot of RNA-sequencing results for Δ*pdhE2* relative to WT. A Log_2_fold > 2 and P-value (Benjamini-Hochberg) < 0.05 were used as significance cutoffs (dashed lines, blue points are significant). *N* = 3 biological replicates. Selecting a functional category on the left-hand panel highlights associated genes in the volcano plot.

**Interactive Plot 6.** Interactive volcano plot of RNA-sequencing results for Δ*manA* relative to WT. A Log_2_fold > 2 and P-value (Benjamini-Hochberg) < 0.05 were used as significance cutoffs (dashed lines, blue points are significant). *N* = 3 biological replicates. Selecting a functional category on the left-hand panel highlights associated genes in the volcano plot.

## Notes

### Competing Interest Statement

The authors have declared no competing interest.

